# Second Generation Drosophila Chemical Tags: Sensitivity, Versatility and Speed

**DOI:** 10.1101/095398

**Authors:** Ben Sutcliffe, Julian Ng, Thomas O. Auer, Mathias Pasche, Richard Benton, Gregory S. X. E. Jefferis, Sebastian Cachero

**Author notes:** These authors contributed equally to this work (Ben Sutcliffe, Sebastian Cachero). Corresponding authors - Gregory Jefferis, Division of Neurobiology, MRC Laboratory of Molecular Biology, Francis Crick Avenue, Cambridge, CB2 OQH, UK. - Sebastian Cachero, Division of Neurobiology, MRC Laboratory of Molecular Biology, Francis Crick Avenue, Cambridge, CB2 OQH, UK.

## Abstract

Labeling and visualizing cells and sub-cellular structures within thick tissues, whole organs and even intact animals is key to studying biological processes. This is particularly true for studies of neural circuits where neurons form sub-micron synapses but have arbors that may span millimeters in length. Traditionally labeling is achieved by immunofluorescence; however diffusion of antibody molecules (>100 kDa) is slow and often results in uneven labeling with very poor penetration into the centre of thick specimens; these limitations can be partially addressed by extending staining protocols to over a week (*Drosophila* brain) and months (mice). Recently we developed an alternative approach using genetically encoded chemical tags CLIP, SNAP, Halo and TMP for tissue labeling; this resulted in >100 fold increase in labeling speed in both mice and *Drosophila*, at the expense of a considerable drop in absolute sensitivity when compared to optimized immunofluorescence staining. We now present a second generation of UAS and LexA responsive CLIP, SNAPf and Halo chemical labeling reagents for flies. These multimerized tags with translational enhancers display up to 64 fold increase in sensitivity over first generation reagents. In addition we developed a suite of conditional reporters (4xSNAPf tag and CLIP-SNAP-Halo) that are activated by the DNA recombinase Bxb1. Our new reporters can be used with weak and strong GAL4 and LexA drivers and enable stochastic, intersectional and multicolor Brainbow labeling. These improvements in sensitivity and experimental versatility, while still retaining the substantial speed advantage that is a signature of chemical labeling, should significantly increase the scope of this technology.

## 1 Introduction

Visualizing molecules in intact tissues with high sensitivity and specificity is of paramount importance in many fields of biological research. Traditionally cellular and subcellular labeling has depended on immunostaining that combines primary antibodies specific to a molecule of interest, followed by labeled secondary antibodies. Recently we and others have adapted chemical labeling approaches that were initially developed for in *vitro* or single cell studies (Keppler et al., 2003; Gautier et al., 2008; Los et al., 2008) for use in genetically defined cells within intact fly and mouse tissues (Kohl et al., 2014; Yang et al., 2015). These overcame a fundamental limitation of antibodies: low diffusion rate that causes poor penetration of thick tissue samples. The basic principle of chemical labeling is the use of small protein tags (engineered from enzymes) that can covalently and irreversibly bind small molecule substrates. These substrates can be conjugated with a variety of labels such as fluorophores for light microscopy and colloidal gold for electron microscopy (Keppler et al., 2003; Gautier et al., 2008; Vistain et al., 2016). High efficiency binding in combination with small substrate size allows easy tissue penetration and fast quantitative staining (Kohl et al., 2014).

Improvements in speed and penetration achieved by the first generation of chemical labeling reagents are particularly important in neural circuit research where labeling of neurons in deep structures within intact brains is essential for understanding connected networks in the brain but experimentally very challenging. To illustrate this point, optimal immunostaining of a fly brain takes more than a week (Ostrovsky et al.,2013) while a mouse brain can take months even when combined with tissue clearing methods (Chung et al., 2013). In contrast, multicolor chemical labeling of a fly brain can be completed within 1 hour, with less than 10 minutes of staining time. Other important advantages of chemical labeling are that it reduces off-target labeling and as completely synthetic reagents, in contrast to antibodies, they are not produced using animals. In comparison to the use of genetically encoded fluorescent proteins, reporter lines with chemical labeling transgenes enable rapid testing and switching to new fluorophores with properties required for constantly evolving imaging modalities. While the published *Drosophila* reagents offer unparalleled staining speed (Kohl et al.,2014), they produce considerably weaker signal than traditional immunolabeling of genetically encoded reporters, limiting their use to relatively strong Gal4 driver lines (Brand and Perrimon, 1993). We now introduce a second generation of fly reagents with greatly increased sensitivity. Furthermore, we have increased the versatility of the system by developing reporters for the LexA-based expression system (Lai and Lee, 2006) and reagents for conditional and stochastic labeling based on Bxb1 DNA recombinase (Huang et al., 2011). Finally we show the utility of chemical labeling in targeting challenging tissues such as the fly antennae. We expect these new tools will greatly increase the use of chemical labeling within the research community, especially speeding up projects that require large numbers of stainings.

## 2 Materials and Methods

### 2.1 Drosophila stocks

Fly stocks were maintained at 25° on iberian food. The driver lines used in this study are MZ19-Gal4 (Ito et al., 1998), MB247-Gal4 (FlyBaseID: FBst0050742), Fru-Gal4 (gift from Barry Dickson) (Stockinger et al., 2005), BG57-Gal4 (FlyBaseID: FBst0032556), GMR50A02-Gal4 (FlyBaseID: FBti0136386), GMR54F05-Gal4 (FlyBaseID: FBst0039080), GMR59F02-Gal4 (FlyBaseID: FBst0039221), OR22a-Gal4 (gift from Leslie Vossall lab) (Vosshall et al., 2000), IR84a-Gal4 (gift from Richard Benton) (Silbering et al., 2011), Orco-LexA::VP16 (gift from Tzumin Lee) (Lai and Lee, 2006), GH146-LexA::GAD (gift from Tzumin Lee) (Lai et al., 2008), nSyb-LexA::P65 in attP40 (Pfeiffer et al., 2012), MB247-LexA (Pitman et al., 2011). The reporter lines used in this study are UAS-CD4::CLIPf on 2nd and 3rd, UAS-myr::SNAPf in attP40 and attP2, UAS-myr::Halo2 in attP40 (Kohl et al., 2014), for details of the new reporter lines generated in this study see Table S1. All images are of female brains, apart from the brains in Figure 4d which are male, all flies were dissected 3-4 days after eclosion.

### 2.2 Drosophila constructs and transgenic flies

Drosophila transformation plasmids from Table 1 were made by Gibson assembly (Gibson et al., 2009) (Figures S4 to S13) or restriction enzyme cloning (Figures S14 to S19). Figures S4 to S19 show the primers and enzymes used to make each plasmid. Transgenic flies were made by BestGene.

**Table 1:** Drosophila transformation plasmids.

| Plasmid name | Genebank accession no. | cloning schematic |
| --- | --- | --- |
| UAS-myr::4xCLIPf | to be added | **Figure S6** |
| LexAop2-myr::4xCLIPf | to be added | **Figure S8** |
| UAS-myr::4xSNAPf | to be added | **Figure S7** |
| LexAop2-myr::4xSNAPf | to be added | **Figure S9** |
| UAS-myr::>HA-BxbSTOP>myr::4xSNAPf | to be added | **Figure S10** |
| LexAop2-myr::>HA-BxbSTOP>myr::4xSNAPf | to be added | **Figure S11** |
| HeatShock-Bxb1-SV40 | to be added | **Figure S4** |
| HeatShock-Bxb1 | to be added | **Figure S12** |
| pJFRC-MUH-stop cassette bxbp | to be added | **Figure S13** |
| JFRC-MUH-FRT-bxbp | to be added | **Figure S13** |
| JFRC81-BxbCassette_Clip_Snap_Halo | to be added | **Figure S5** |
| UAS-Halo7::CAAX | to be added | **Figure S14** |
| UAS-3xHalo7::CAAX | to be added | **Figure S15** |
| UAS-7xHalo7::CAAX | to be added | **Figure S16** |
| UAS-Syt::Halo7 | to be added | **Figure S17** |
| UAS-3xSyt::Halo7 | to be added | **Figure S18** |
| UAS-7xSyt::Halo7 | to be added | **Figure S19** |
| UAS-LA::Halo2 | to be added | **Figure S20** |

### 2.3 Labeling Reagents

Substrates were acquired either as stock solutions (e.g., HaloTag-TMR) or in powdered form (SNAPf and CLIPf substrates) and diluted/dissolved in anhydrous dimethyl sulfoxide (DMSO) (Life Technologies) to a concentration of 1 mM. Aliquots (5 μL) were stored at −20° in the presence of desiccant. We observed that using old DMSO or storing dissolved substrates in moist and/or warm conditions can lead to hydrolysis, reducing labeling efficiency. For a list of all substrates used in this study see Table 2.

**Table 2:** Chemical Tagging Substrates used in this study. Commercially available,fluorophore-coupled substrates for SNAP-, CLIP- and Halo- are listed.

| Substrate<br>(abbreviation) | Fluorophore | Ex | Em | Binds to | Cell<br>permeable | Supplier | Cat.<br># |
| --- | --- | --- | --- | --- | --- | --- | --- |
| SNAP-Cell 647-SiR<br>(SNAP-SiR) | SiR | 645 | 661 | SNAPm/f | Yes | NEB | S9102S |
| SNAP-Surface 549<br>(SNAP-549) | Dyomics<br>DY-549P1 | 560 | 575 | SNAPm/f | No | NEB | S9112S |
| CLIP-Surface 488<br>(CLIP-488) | ATTO-TEC<br>488 | 506 | 526 | CLIPm/f | No | NEB | S9232S |
| CLIP-Surface 547<br>(CLIP-547) | Dyomics<br>DY-547 | 554 | 568 | CLIPm/f | No | NEB | S9233S |
| HaloTag TMR Ligand<br>(Halo-TMR) | TMR | 555 | 585 | Halo2/7 | Yes | Promega | G8252 |
| HaloTag SiR Ligand<br>(Halo-SiR) | SiR | 645 | 661 | Halo2/7 | Yes | K. Johnsson | n/a |

### 2.4 Protocol for labeling Drosophila Brains

Single and double channel labeling of Drosophila brains was carried out as previously described (Kohl et al., 2014). For labeling of UAS-LA::Halo2 fillet preparation of wandering third instar larvae were made followed by the same protocol used for labeling whole brains. For detailed information on staining Chemical Brainbow brains and antennal segments see Supplemental Information. We find that CLIPf substates weakly bind SNAPf tag, therefore if labeling both SNAPf and CLIPf in the same specimen we recommend doing sequential SNAPf substrate incubation (minimum 5 min) then addition of CLIPf substrate (minimum 5 min), to avoid cross reactivity.

### 2.5 Image Acquisition and Deconvolution

Confocal stacks of fly brains were imaged at 768 × 768 pixels every 1 μm (voxel size of 0.46 × 0.46 × 1 μm; 0.6 zoom factor) using an EC Plan-Neofluar 40 ×/1.30 Oil DIC M27 objective and 16-bit color depth. Higher magnification images of cell bodies were acquired at 2048 × 2048 pixels every 0.45 μm (voxel size 0.1 × 0.1 × 0.45 μm; 1.0 zoom factor) using a Plan-Apochromat 63×/1.40 Oil DIC M27 objective and 16-bit color depths. Antennae were imaged at 1024 × 1024 pixels every 1 μm (voxel size 0.20 × 0.20 × 1 μm; 1.0 zoom factor) using an EC Plan-Neofluar 40x/1.30 Oil DIC M27 objective and 8-bit color depths. The image of the entire larval musculature (Figure 5b) was acquired as a tile scan with total dimensions 1536 × 2304 pixels every 1.0 μm (voxel size 1.84 × 1.84 × 1.0 μm; 0.6 zoom factor) with EC Plan-Neofluar 10×/0.30 M27 objective and 16-bit color depths. The high magnification larval muscle inset was acquired at 2156 × 2156 pixels every 0.45 μm (voxel size 0.1 × 0.1 × 0.45 μm; 1.0 zoom factor) using a Plan-Apochromat 63×/1.40 Oil DIC M27 objective and 16-bit color depth. All images acquired on a Zeiss LSM710 confocal microscope.

The confocal stack of the fly brain in Figure 1d was acquired using a Leica SP8 confocal microscope, following the Nyquist criterion, at 4224 × 4224 pixels every 0.313 μm (voxel size 0.076 × 0.076 × 0.313 μm; 0.9 zoom factor) using a HC PL APO CS2 40×/1.30 oil objective. Image deconvolution was carried out on each channel individually using the Huygens Professional (Scientific Volume Imaging) software with a backprojected pinhole of half the emission wavelength in nm, a theoretical Point Spread Function, automatic back ground estimation, 5 Iterations, a Signal to noise ration of 20, a Quality threshold of 0.05, optimized Iteration mode and an automatic brick layout. The separate deconvolved channels were then combined as an RGB tiff using Fiji (Schindelin et al., 2012).

**Figure 1:**
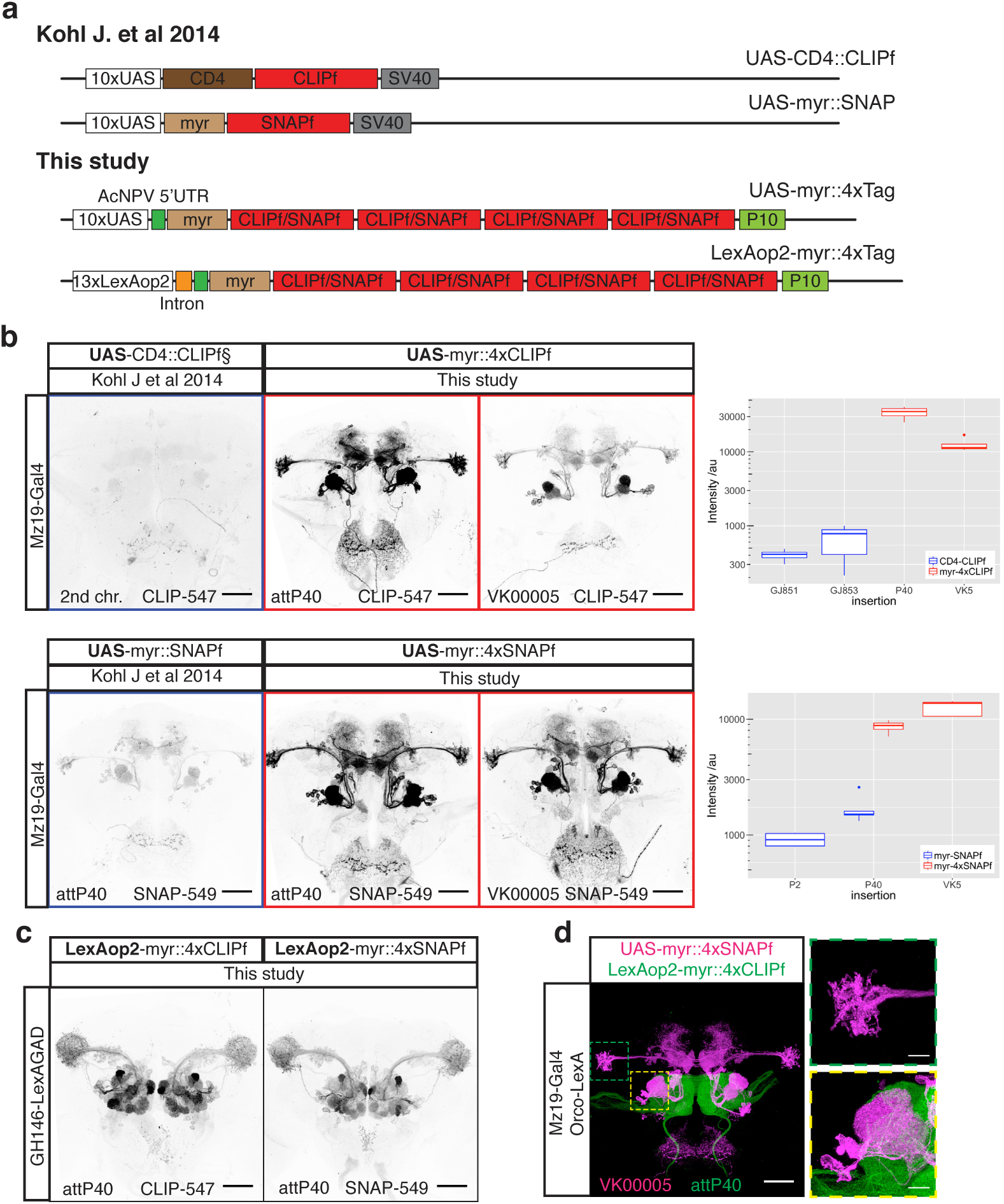
New CLIPf and SNAPf reporters have increased sensitivity. **(a)** Schematic of previous CLIPf/SNAPf reporters from (Kohl et al., 2014) and the new reporters from this study. **(b)** Labeling of Mz19-Gal4 neurons using the old and new reporters. Each panel contains information on the dye used and insertion sites. Box plots show the quantification of fluorescence intensity of the axonal terminals of projection neurons in the lateral horn (arbitrary units). Boxplot n numbers were; GJ853 CD4::CLIPf on 2nd n=3, GJ851 CD4::CLIPf on 3rd n=4, P40 myr::4xCLIPf n=4, VK00005 myr::4xCLIPf n=4, P40 myr::4xSNAPf n=4, VK00005 myr::4xSNAPf n=5, P2 myr::SNAPf n=4 and P40 myr::SNAPf n=5 **(c)** New LexAop2-myr::4xCLIPf/4xSNAPf reporters labeling olfactory projection neuron using the weak GH146-LexA::GAD driver. **(d)** Orthogonal labeling of olfactory sensory neurons (green) and projection neurons (magenta) using new tags. Shown is the max intensity projection of a confocal stack after deconvolution. Images in panels b and c were acquired using the same microscope settings. Scale bars are 50 μm in whole brain images and 10 μm in higher magnification images of the boxed areas in panel d.

### 2.6 Fluorescence quantification

For the comparison between old and new reporters we acquired confocal stacks using two different 561 nm laser power settings (low 2% and high 10%) with gain (600) and pinhole (60.1 μm, 1.42 AU) remaining constant. Images acquired at the low setting were optimal for non-saturated images of the new reporters and images acquired at the high setting were optimal for the old reporters so that we had a stack that could be segmented for quantification and then the data from the low stacks were quantified (see below). Confocal .lsm files were then converted to .nrrd files using Fiji. Using Amira 6.0.1 (FEI, Thermo Fisher Scientific) a .nrrd stack, for each brain to be quantified, was opened (high versions for the old reported and low versions for the new reporters) and a median filter of 3 iterations was applied. Using the Segmentation Editor in Amira 6.0.1, two materials were assigned to the median filtered stack for each brain: 1) for quantifying signal a three dimensional ROIs surrounding the axonal terminals of Mz19-Gal4 PNs in the lateral horn, 2) for background correction a three dimensional region ventral to the axonal terminals of Mz19-Gal4 PNs in the lateral horn.The intensity and background correction calculations were performed in R (Team, 2016) and detailed in R Markdown supplemental file. Briefly, for comparison of the old and new CLIPf reporters we used the average intensity in the LH of the old reporters as baseline and then divided the quantified intensity of the new reporter by the average for the old reporters to give a fold change (e.g. for the comparison of new 4xCLIPf in attP40 with the old version of the CLIPf reporters: the intensity value of 4xCLIPf in attP40 was divided by the average of the intensities calculated for both insertions of the old version CLIPf reporters, see the R Markdown supplemental file for details of the calculations). For new vs old comparisons of the Halo reporters we calculated percentage change as this was a more meaningful comparison (see the R Markdown supplemental file for details of the calculations).

## 3 Data Availability

All data necessary for confirming the conclusions presented in the article are represented fully within the article. All fly strains and plasmids are available upon request. Sequence data for all plasmids will be made available at GenBank and the accession numbers listed in Table 1. Code used to quantify fluorescence intensities is provided in File S1.

## 4 Results

### 4.1 New CLIPf and SNAPf reporters with increased sensitivity

The first generation of chemical labeling reporters achieved rapid staining times, shortening protocols from over 100 hours to less than 1 hour for whole mount *Drosophila* brains (Kohl et al., 2014). Despite this dramatic improvement in staining speed, signal strength is lower than antibody staining of reporter proteins. This is likely due to the non-amplifiying nature of chemical labeling: one molecule of tag covalently binds one substrate molecule fused to one molecule of fluorophore. This linearity can be beneficial when quantifying signal intensity. In contrast, with immunofluorescence one target can be bound by more than one primary antibody which is then recognized by several secondary antibody molecules each conjugated to multiple fluorophores leading to substantial signal amplification. This lower sensitivity is evident when comparing the signal from several Gal4 lines (Rubin collection, Janelia Research Campus) driving GFP or first generation CLIPf and SNAPf reporters (Figures S1a and S2). To bridge this gap and extend the use of chemical labeling to most Gal4 driver lines, weak and strong, we designed a new generation of reporters with greatly increased sensitivity. These reporters differ from the original ones in two ways: first, they have a short 5’ UTR (AcNPV) and the 3’ UTR from the *A. californica nucleopolyhedrovirus* P10 gene; these modifications have been shown to increase translational efficiency by more than 20 times (Pfeiffer et al., 2012) and second, they are tetramerized to increase reporter signal up to four fold (Shearin et al., 2014) (Figure 1a). We generated transgenic fly lines by inserting these new 4xCLIPf and 4xSNAPf reporters into the well-characterized attP40 and VK00005 phiC31 landing sites on the 2nd and 3rd chromosomes, respectively (Table S1).

We tested these new transgenes and compared them to the first generation reporters using the sparse line Mz19-Gal4, a driver of medium strength that expresses in about 12 olfactory projection neurons innervating three adjacent olfactory glomeruli and a group of neurons with processes near the mushroom bodies. When driven by Mz19-Gal4 all reporters produced the expected labeling pattern. In comparison, the first generation tags were barely visible when imaged under conditions that produced strong signal with the new reporters (Figure 1b). To quantify the increase in signal strength we measured intensity in the axonal terminals of projection neurons in the lateral horn (green dotted area in Figure 1d, see methods). Using the average between UAS-CD4::CLIPf on the 2nd and 3rd chromosomes as baseline, the new UAS-myr::4xCLIPf reporters are 64 (attP40) and 24 (VK00005) times brighter. In the case of SNAP, the new UAS-myr::4xSNAPf reporters are 7 (attP40) and 10 (VK00005) times brighter than the average between the first generation UAS-myr::SNAPf in attP2 and attP40. While CLIPf and SNAPf substrates use different fluorophores and have different labeling sensitivities, complicating precise quantitative comparisons, the new CLIPf and SNAPf reporters produced qualitatively similar fluorescence intensities. To extend these results to other driver lines we used a number of Gal4 P element and enhancer fusion insertions of varying strengths to drive the new reporters (weakest to strongest: GMR-50A02-Gal4, GMR-59F02-Gal4 and GMR-54F05-Gal4). Qualitatively these stainings recapitulated the Mz19-Gal4 results with the new reporters showing large increases in brightness (Figure S1 and S2). These results indicate the new reporters are suitable for labeling most if not all Gal4 driver lines that show expression after immunostaining.

### 4.2 LexA responsive reporters

Dissecting the function of neuronal components in a circuit often requires labeling more than one cell population with different reporters that respond to orthogonal drivers such as Gal4 and LexA. To increase the flexibility of the chemical labeling platform we made LexA responsive tetramerized CLIPf and SNAPf reporters and inserted them in attP40 and VK00005 (Table S1). We tested these reporters using the weak driver line GH146-LexA::GAD. We found that LexAop2-myr::4xCLIPf and LexAop2-myr::4xSNAPf reporters inserted in both chromosomal locations produced strong labeling (Figure 1c and Figure S2c). Since new LexA drivers are now routinely made with the strong p65 transactivation domain rather than the weaker GAD domain, this result suggests our new reporters will be useful for most LexA driver lines. Finally, we show how these new reagents can be used for visualizing different cell populations by labeling olfactory sensory neurons (Orco-LexA::VP16) and a subset of their post-synaptic projection neurons (Mz19-Gal4) in the same brain (Figure 1d). While we imaged this brain using a confocal microscope (following the Nyquist criterion and subsequent deconvolution, see methods), super-resolution microscopy techniques, such as STED, could also be used, when available for thick tissue specimens, to increase resolution.

### 4.3 New Halo tag reporters with improved membrane localization and signal strength

Our first generation Halo tag reporters already incorporated the 5’ and 3’ translational enhancers L21 and P10 (Figure 2a) and were inserted into PhiC31 landing sites that support strong expression (attP40 and attP2). While this tag produced the brightest signal among the first generation of chemical reporters we noticed an unexpected accumulation of the tag in the cell nucleus and reduced signal in axons (Figure 2b) suggesting suboptimal cellular localization. Intriguingly, 4xCLIPf and 4xSNAPf tags use the same myristoylation signal as Halo (first 90 amino acids from the *Drosophila* Src protein) but are excluded from the nucleus, displaying the expected membrane localization. In order to improve cellular localization we replaced the N-terminal myristolation with a C-terminal CAAX membrane targeting signal (Choy et al., 1999). In addition we made several reporters with either one, three and seven tandem fusion-tags of Halo with the aim of increasing labeling efficiency (Figure 2a). The new constructs use Halo version 7 (Halo7) which is reported to show increased expression, stability and substrate binding kinetics over version 2 (Halo2) (Encell et al., 2012). We made transgenic flies with insertions in attP40, VK00005 and VK00027 (Table S1).

**Figure 2:**
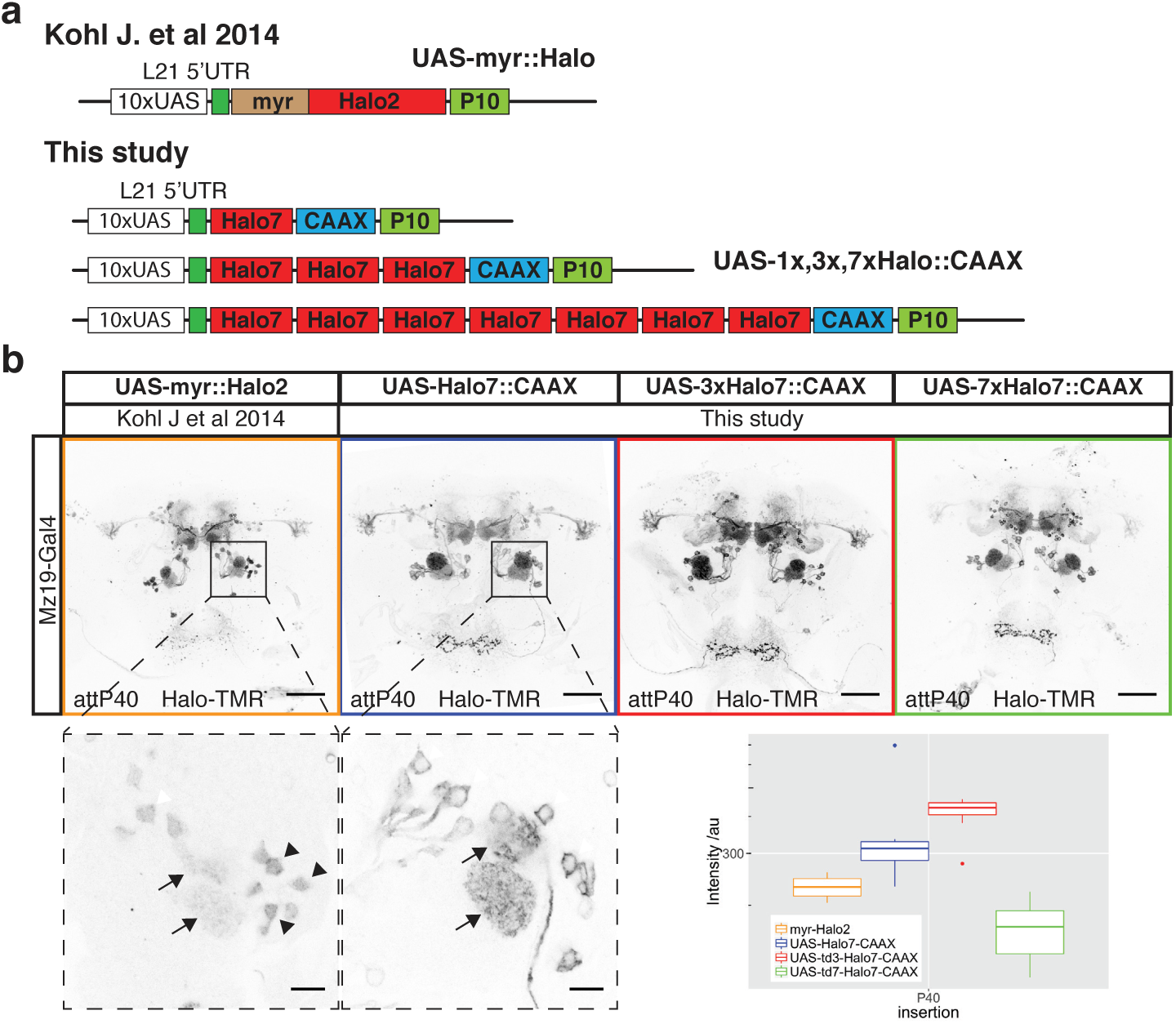
New Halo reporters with improved sensitivity and localization. (**a**) Schematic of CLIP/SNAPf reporters from (Kohl et al., 2014) and the new reporters from this study. **(b)** Labeling of Mz19-Gal4 positive neurons using the old myr::Halo2 and new Halo7::CAAX reporters. All images were aquired using the same microscope settings. Lower panels are high magnification single slice images showing differences in reporter localization in the cell bodies (arrowheads) of olfactory projection neurons. Arrows indicate signal in glomeruli. The box plot shows the quantification of fluorescence intensity of the axonal terminals of PNs in the lateral horn (arbitrary units). Boxplot n numbers were; myr::Halo2 n=7, UAS-Halo7::CAAX-P40 n=7, UAS-3xHalo7::CAAX n=8 and UAS-7xHalo7::CAAX n=8. Scale bars in full brain images are 50 μm and higher magnification images of cell bodies 10 μm.

We compared cellular localization and signal intensities from the first and new generation of Halo tags in the same way as for CLIPf and SNAP. Nuclear signal is greatly reduced in the new CAAX reporters when compared to the myristoylated ones (See higher magnification images from the first two panels of Figure 2b). In addition, we measured modest increases in signal strength with the new monomeric and trimeric reporters (53% and 78% brighter, respectively, Figure 2b, box plot). Surprisingly the heptamer is 28% less bright than the old reporter, possibly due to increased instability or impaired trafficking (Figure 2b, box plot).

### 4.4 Chemical tags in peripheral sensory organs

We wanted to explore the performance of chemical labeling in tissues other than the brain, where differences in extracellular matrix or other cellular barriers may have a negative impact on labeling. To accomplish this we stained sensory neurons in whole-mount third antennal segments. This tissue is typically regarded as hard to stain in part because it is surrounded by cuticle, in contrast to brains which are dissected out of the head capsule before staining. While immunolabeling can work, as for brains, the optimized protocol spans up to a week (Saina and Benton, 2013). Using GAL4 driver lines that label sensory neurons (Ionotropic receptor 84a (IR84a) and Odorant receptor 22a (Or22a)), we expressed the new 4xSNAPf and 3xHalo7 reporter lines in the antennae (Figure 3 and S3). While reporters produced signal in the expected cells in all cases, shorter labeling incubations produce lower background, especially in the cuticle (Figure 3b, arrowheads). The SNAPf label also resulted in more uniform labeling of the axons and soma when compared to a mCD8::GFP reporter (Figure 3a, arrowheads vs arrows). In contrast to immunostaining, chemical labeling reagents penetrate rapidly as demonstrated by the signal being as strong after 10 minutes as it is after 3 h (Figure 3b). In addition chemical labeling in the antennae, as in the brain (Kohl et al., 2014), can be combined with immunolabeling, in this case of the Or22a receptor (Figure S3).

**Figure 3:**
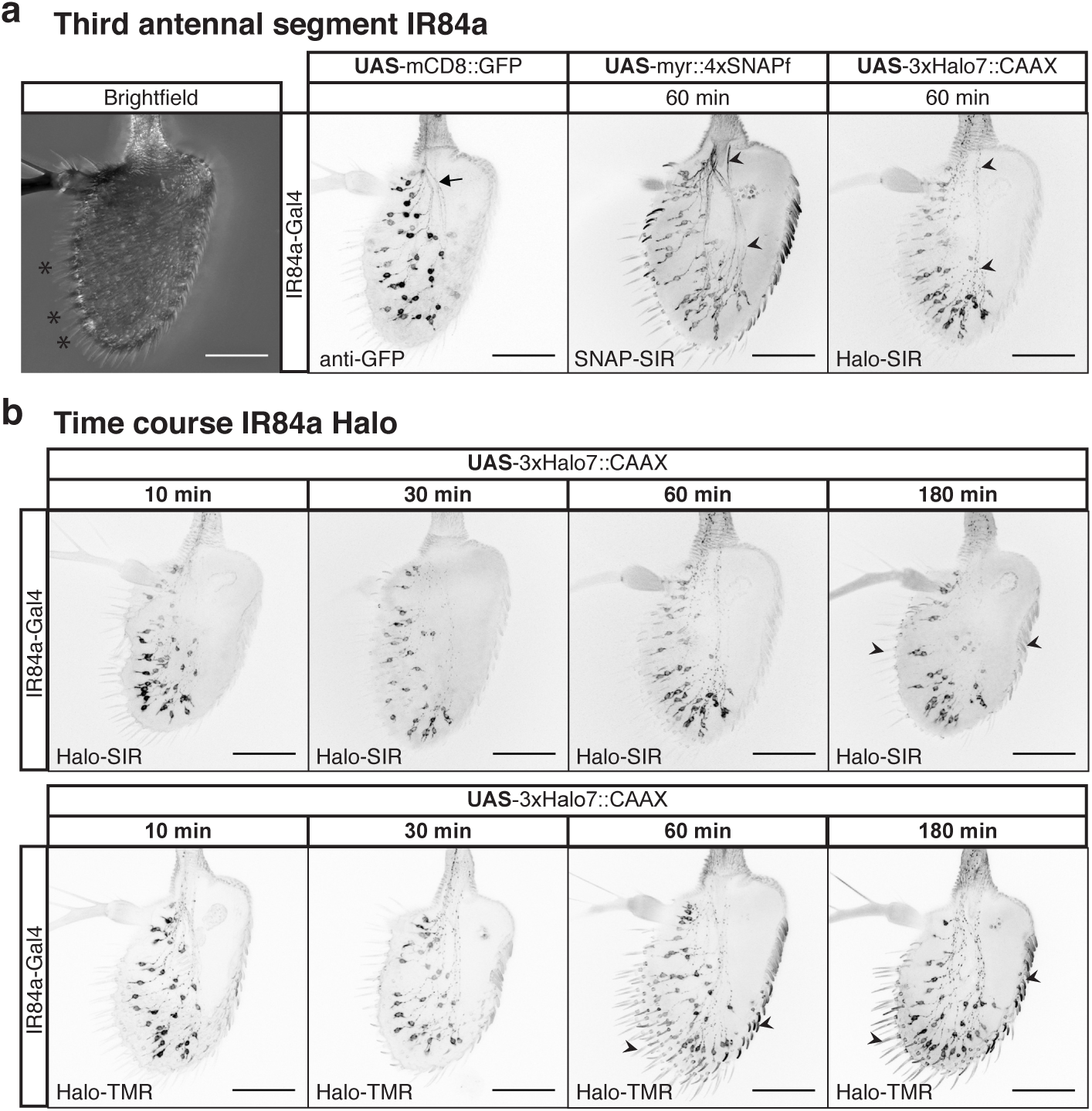
Chemical tags in peripheral sensory organs. (a) Left-most image, bright field image of the antennae, sensilla are marked with asterisks. Chemical labeling of Ionotropic Receptor 84a (IR84a) expressing sensory neurons. Comparison between GFP immunostaining and SNAP-SiR chemical labeling; arrow and arrowheads highlight stronger labeling of axons by chemical labeling relative to the soma. **(b)** Incubation time series for far red Halo-SiR (top row) and red Halo-TMR (bottom row) dyes. All panels shows partial projections of confocal stacks that exclude the cuticle. Scale bars are 50 μm.

### 4.5 Conditional reporters

Brains are densely packed with neurons of great diversity both in morphology and function. A fundamental step in studying complex neural circuits is to break them down into smaller components by visualizing the morphology of single or small clusters of neurons. To achieve this, neuroanatomical studies take advantage of large promoter-Gal4 and LexA collections to find sparse drivers to express reporters in small numbers of neurons. While this approach greatly limits the number of labeled cells, they often have overlapping processes which cannot be resolved by light microscopy. In these cases further labeling refinements, using a number of genetic strategies, are often required (Jefferis and Livet, 2012). We extended the applicability of chemical labeling to these situations by developing reagents to: a) limit the number of labeled cells or b) increase the combinatorial number of fluorophores available for each labelled neuron.

To limit the number of labeled cells we designed an inactive reporter with a transcriptional stop cassette upstream of the coding region for 4xSNAP. This reporter can be activated upon removal of the stop cassette by the DNA recombinase Bxb1 (Figure 4a). We chose Bxb1 from mycobacteriophage (Huang et al., 2011) as it is orthogonal to recombinases commonly used in *Drosophila.* Another advantage is its irreversibility as it recombines attP and attB sites to generate new attL and attR sites which are no longer substrates. We generated lines that express Bxb1 in three different ways: a) stochastically, using a heat shock inducible promoter (hs-Bxb1, Figures S4 and S12), b) by driving its expression with Gal4 (UAS-Bxb1, Figure 4b) and c) by using a combination of Gal4 and Flp DNA recombinase (UAS>FlpSTOP>Bxb1, Table S1).

**Figure 4:**
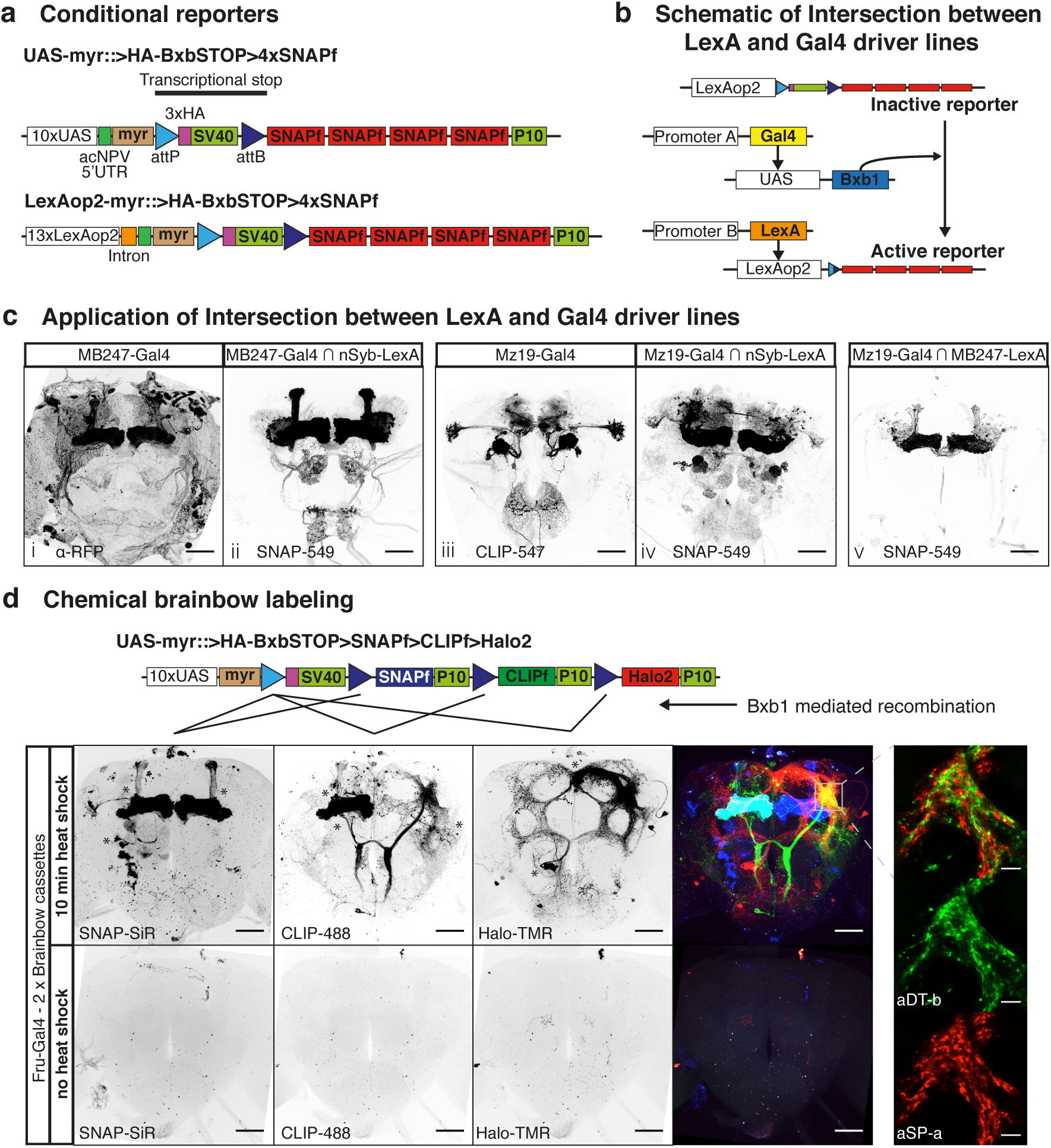
Sparsening expression using conditional chemical reporters. **(a)** Schematic of new conditional reporters. The HA tag present in the stop cassette can reveal the expression of the inactive reporter (not shown). **(b)** Schematic showing the genetic approach to intersect LexA and Gal4 in c. **(c)** Panels i and iii show confocal projections of Gal4 lines driving regular reporters. Panels ii and iv show confocal projections of Gal4 lines intersected with the panneuronal nSyb-LexA::P65 using the scheme from b. Panel v shows a confocal projection of the intersections between the sparse lines Mz19-Gal4 and MB247-LexA::VP16. **(d)** Heatshock activation of Brainbow cassettes during early development label neuroblast clones of *fruitless* positive neurons. Bottom panels show the Brainbow cassettes are silent when no heatshock is applied. Asterix indicate the cell bodies form neuroblast clones. Panels on the right: high magnification single confocal slice showing the close apposition between processes from the two sexually dimorphic clones aSP-a and aDT-b. Scale bars for full brain images are 50 μm and scale bars for higher magnifications are 10 μm.

As a proof of principle we used the conditional reporters in three experiments to intersect the expression of Gal4 and LexA drivers. The schematic in Figure 4b shows the logic of the experiment: MB247-Gal4 or Mz19-Gal4 drives expression of UAS-Bxb1 to activate the conditional reporter LexAop2-myr::>BxbSTOP>4xSNAP; the activated reporter is then driven by MB247-LexA::VP16 or the pan-neuronal nSyb-LexA::p65. In the first experiment, MB247-Gal4 ∩ nSyb-LexA::P65, the result is very similar to that of a regular reporter with the exception of the lack of strong glial staining, normally present in MB247-Gal4, due to the reporter being driven by the neuronal specific nSyb-LexA::p65 (compare Figure 4c.i and c.ii). On the other hand, the second experiment shows Mz19-Gal4 ∩ nSyb-LexA::P65 is considerably broader than that of the regular reporter including labeling in the mushroom bodies (compare Figure 4c.iii and c.iv). Mz19-Gal4 ∩ nSyb-LexA::P65 reflects two interesting properties of this approach: first, it captures and immortalizes developmental expression and second, weakly expressing cells, previously undetectable with a regular reporter, could drive Bxb1 mediated recombination allowing strong reporter expression driven by nSyb-LexA::P65. In the third experiment we used Mz19-Gal4 to activate the reporter and MB247-LexA::VP16 to drive it; as one would predict from the previous two experiments this intersection labels a modest number of mushroom body Kenyon cells (Figure 4c.v).

The second strategy for resolving overlapping processes is multiplexing the label. The approach we took is based on the Brainbow technique (Livet et al., 2007; Hadjieconomou et al., 2011; Hampel et al., 2011) using the tags CLIPf, SNAPf and Halo2 (Figure 4d). Our reporter incorporates translational enhancers without multimerization. We used Bxb1 to activate the cassette as for our single tag conditional reporters. Because Bxb1 recombination is irreversible the cassette requires fewer recombination sites than previous Brainbow reporters. Upon expression of the recombinase, the single attP site recombines with one of the three attB sites removing the intervening DNA and irreversibly selecting one of the three tags for expression (see schematic in Figure 4d). We made fly lines with the Brainbow cassette inserted into attP2 and VK00005 (Table S1).

We tested the new cassettes by labeling subsets of neurons that express the male specific form of the Fruitless protein (FruM). By activating the Brainbow cassette immediately after larval hatching we aimed to create groups of labeled cells of the same developmental origin (neuroblast clones, see methods). Our pilot experiment showed that both transgenes are efficiently activated producing the expected *fruitless* positive neuroblast clones (Compare Figure 4d with Cachero et al. (2010)). We found that the three chemical tags were activated in a similar number of neuroblast clones (marked with asterisks in Figure 4d: 3 clones for SNAPf, 3 for CLIPf 3 and 2 for Halo2). The presence of both Brainbow cassettes can can be seen in the mushroom body clone on the fly’s right side where both CLIPf and SNAPf tags were activated, labeling the resulting clone in cyan. Resolving several clones in a single brain has the advantage of requiring fewer samples to describe the anatomy of a neuronal population. Furthermore it enables researchers to examine the overlap between clones within the same brain rather than using image registration and post hoc comparisons of clones from multiple brains. For instance, it makes possible to examine the close apposition of processes from aSP-a and aDT-b clones in the male enlarged region of the brain (Figure 4d, high magnification insets).

### 4.6 Sub-cellular reporters

Encouraged by the good results obtained while labeling membranes, we wanted to make reporters for other cellular compartments, both in the nervous system and elsewhere.

Synapses are the key sites of information transfer in neuronal circuits. In order to label them we made UAS reporters where one, three or seven copies of the Halo7 tag are fused to the pre-synaptic protein Synaptotagmin (Syt, Table S1). When driven by Mz19-Gal4 all three Syt::Halo7 synaptic markers produced strong labeling in areas known to have presynapses with minimal presence in regions devoid of them (compare Figure 2b and 5a). The gradation in signal strength going from monomer to heptamer makes these reporters useful for labeling synapses using drivers ranging from weak to strong.

Next, we made a reporter for fast and sensitive labeling of actin filaments by fusing a peptide, LifeAct (LA) that binds actin filaments to Halo2 (Table S1) (Riedl et al., 2008). As a proof of principle we expressed the reporter using the pan-muscular driver BG57-Gal4. These larvae are viable despite widespread expression of LA::Halo2, indicating the reporter is not overtly toxic. The staining of body wall muscles in third instar larvae revealed the expected expression pattern with stripes of muscle actin bundles clearly visible (Figure 5b).

**Figure 5:**
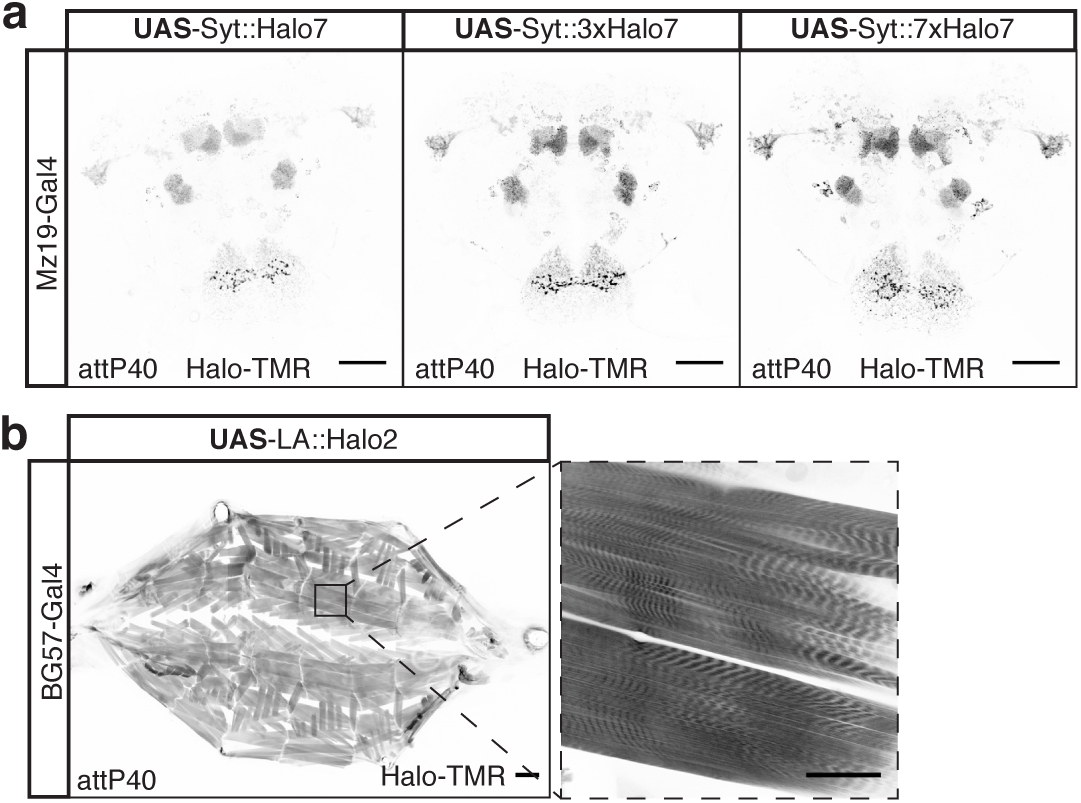
New sub-cellular Halo reporters. **(a)** Labeling the synaptic terminals of Mz19-Gal4 positive neurons using Halo7 reporters fused to Synaptotagmin. **(b)** Labeling of muscle actin in the larva using a fusion between Halo2 and LifeAct peptide. Scale bars in full brain images and higher magnification of muscle fibre are 50 μm and full larva 200 μm.

## 5 Discussion

In this study we introduce a second generation of chemical tags that achieve substantial improvements in sensitivity and versatility over the first generation. Most applications where tag immunostaining is used can benefit from super fast and highly sensitive chemical labeling and the new reagents are ideally suited for medium to high throughput applications such as anatomical screens of driver lines or assessment of RNAi screen phenotypes.

The introduction of LexAop2 and conditional reporters opens the possibility to a larger set of experiments than was possible with first generation reagents. For instance combining UAS and LexAop2 reporters will allow super-resolution microscopy to resolve potential contacts between different neuronal populations. The Brainbow cassette can be used in large anatomical screens enabling rapid characterization of complex driver lines by labeling multiple clones in the same brain (Livet et al., 2007; Hadjieconomou et al., 2011; Hampel et al., 2011). Besides the increase in speed, this allows imaging different neuronal populations in the same brain offering a powerful insight into their potential connectivity. Our conditional reporters can be used to capture developmental expression; these could be exploited for a systematic study of neuronal fate during metamorphosis. While we validated our reagents in the antennae it is likely that chemical labeling will work in most other tissues. Beyond the field of neuroscience the chemical actin reporter will be a useful alternative to the widely used but highly toxic phalloidin staining, particularly in those applications where genetically targeting to specific muscles could be an advantage. A second advantage is the irreversible nature of the chemical staining, while phalloidin stainings fade with time. Lastly, it could be used for *in-vivo* imaging when combined with cell permeable substrates.

The improvements in signal strength achieved by the new reagents derive from their higher expression levels. For experiments where an even stronger signal is needed more than one transgene could be used. In the case of the Brainbow cassettes we are currently multimerizing the tags to obtain higher signal to noise ratio. Another possibility would be developing brighter ligands, for instance by conjugating multiple fluorophores per ligand molecule. The collection of reagents presented here is by no means exhaustive, further additions to this toolkit could include generation of reporters to harness the QUAS system (Potter et al., 2010) and expansion of the multimerized chemical tags to target sub-cellular compartments and organelles; for example axons, dendrites, microtubules and mitochondria.

While the new chemical tags were successful in producing strong labeling of all Gal4 and LexA lines tested, a new comparison between chemical labeling and smFP immunolabeling (Viswanathan et al., 2015) found that the latter still yields better signal to noise ratio than a single copy multimerized chemical tag (Meissner G., personal communication). This is unsurprising as the smFPs are one of the most optimized tags available for immunostaining with 10-15 copies of their epitope tags which are then subjected to a highly optimized, but long (>10 days), staining protocol. Therefore, in our view the significant increase in speed and reproducibility derived from the simple chemical labeling protocol, coupled with strong signal make it an attractive option for most applications.

In conclusion, the new reagents generated in this study significantly extend the experimental reach of chemical labeling to most forms of genetic labeling scenarios in *Drosophila.* This should significantly increase its use by the research community. We hope that this will also encourage non-*Drosophila* researchers to expand and optimize the use of chemical labeling in other genetic model organisms.

## 6 Acknowledgements

We thank members of the G.S.X.E.J. lab for comments on the manuscript. This work was supported by the Medical Research Council [MRC file reference U105188491 and U105178788], European Research Council Starting Investigator (211089) and Consolidator Grants (649111) to G.S.X.E.J and a Royal Society Dorothy Hodgkin Fellowship to S.C.. T.O.A. is supported by a Human Frontier Science Program Long Term Fellowship. Research in R.B.’s laboratory is supported by the University of Lausanne and an ERC Consolidator Grant (615094). Stocks obtained from the Bloomington Drosophila Stock Center (NIH P40OD018537) were used in this study.

## 8 Supplemental Information

### 8.1 Materials and Methods

Labeling of tissues was performed as follows.

### 8.1.1 Triple labeling of Chemical Brainbow Drosophila Brains

We made a double cassette reporter by combing both Brainbow insertions with hs-Bxb1 and *fruitless*-Gal4; a knock in insertion of Gal4 into the P1 promoter of the Fru locus (Stockinger et al., 2005). First we screened several hs-Bxb1 insertions and identified one that produces minimal background activation of the reporters at 25° (that is the line that gave fewest labelled cells with no heat shock). Next, newly hatched larvae were heat-shocked for 10 minutes at 37° and allowed to develop into adults. Flies were then processed as follows:

- All steps were carried out at room temperature unless stated differently.
- Brains were dissected in ice-cold 0.1 M phosphate buffer (PB: 0.032 M NaH_2_PO_4_, 0.068 M NaH_2_PO_4_).
- Fixed in 4 % paraformaldehyde (PFA) (in 0.1 M PB) for 30 min in a glass-well plate on an orbital shaker
- Transferred to 1.5 ml tube
- Permeabilized by incubation in 1 ml of PBS-T (phosphate buffered saline + 0.3 % Triton X-100) (2 × 15 min) on rotating wheel
- Incubated with SNAP-Cell 647-SiR (NEB: S9102S) substrate at a final concentration of 1 μM in PBS on rotating wheel for 30 min
- Added CLIP-Surface 488 (NEB: S9232S) and HaloTag TMR (Promega: G8252) at a final concentration of 1 μM to the tube with brains/SNAP-Cell 647-SiR solution
- Incubated on rotating wheel for a further 30 mins
- Washed with PBS-T (2 × 15 min)
- PBS-T was removed as completely as possible and 100 μl of Vectashield (or other) mounting medium added. We observed that subsequently transferring brains into a fresh 100 μl aliquot of Vectashield results in more homogeneous signal along the z axis of the image
- After labeling brains were then mounted on charged slides and imaged

### 8.1.2 Chemical labeling of Drosophila Antennal Segments

- All steps were carried out at room temperature unless stated differently.
- Antennae were harvested in liquid nitrogen Saina and Benton (2013)
- Fixed in 4% PFA, 3% Triton, 1xPBS (180 min)
- Washed in 3% Triton, 1xPBS (2 × 5 min)
- Washed in 0.1% Triton, 1xPBS (2 × 5 min)
- Incubated in 5 μM Halo-SIR (10 min)
- Washed in 0.1% Triton, 1xPBS (2 × 10min)
- Washed in 0.1% Triton, 1xPBS (2 × 5min)
- After labeling antennae were mounted in Vectashield and imaged

### 8.1.3 Chemical and Antibody co-labeling Drosophila Antennal Segments

- All steps were carried out at room temperature unless stated differently.
- Antennae were harvested in liquid nitrogen
- Fixed in 4% PFA, 3% Triton, 1xPBS (180 min)
- Washed in 3% Triton, 1xPBS (2 × 10min)
- Washed in 0.1% Triton, 1xPBS (2 × 10min)
- Incubated in 5 μM Halo-SIR (10 min)
- Washed in 0.1% Triton, 1xPBS (2 × 10min)
- Blocked in 5% goat serum, 0.1%Triton, 1xPBS (60 min)
- Incubated with primary antibody in in 5% goat serum, 0.1%Triton, 1xPBS (overnight at 4°)
- Washed in 0.1% Triton, 1xPBS (6 × 15min)
- Blocked in 5% goat serum, 0.1%Triton, 1xPBS (60 min)
- Incubated with secondary antibody in in 5% goat serum, 0.1%Triton, 1xPBS (overnight at 4°)
- Washed in 0.1% Triton, 1xPBS (6 × 15min)
- After labeling antennae were mounted in Vectashield and imaged

**Table S1:** Transgenic flies generated in this study

| Tag | Promoter | Insertion site | Condi-tional | Transgene | Figure |
| --- | --- | --- | --- | --- | --- |
| CLIP | UAS | VK00005 | - | UAS-myr::4xCLIPf | 1, S1 |
|  |  | attP40 | - | UAS-myr::4xCLIPf | 1, S1 |
|  | LexAop2 | VK00005 | - | LexAop2-myr::4xCLIPf | S2 |
|  |  | attP40 | - | LexAop2-myr::4xCLIPf | 1 |
| SNAP | UAS | VK00005 | - | UAS-myr::4xSNAPf | 1, S2 |
|  |  | attP40 | - | UAS-myr::4xSNAPf | 1, S2 |
|  | LexAop2 | VK00005 | - | LexAop2-myr::4xSNAPf | S2 |
|  |  | attP40 | - | LexAop2-myr::4xSNAPf | 1 |
|  | UAS | VK00005 | yes | UAS-myr::>BxbSTOP<br>>-4xSNAPf | - |
|  | LexAop2 | VK00018 | yes | LexAop2-myr::>BxbSTOP<br>>-4xSNAPf | 4 |
| Halo | UAS | attP40 | - | UAS-Halo7::CAAX | 2 |
|  |  | attP40 | - | UAS-3xHalo7::CAAX | 2,3 |
|  |  | attP40 | - | UAS-7xHalo7::CAAX | 2 |
|  |  | attP40 | - | UAS-Syt::Halo7 | 5 |
|  |  | attP40 | - | UAS-Syt::3xHalo7 | 5 |
|  |  | attP40 | - | UAS-Syt::7xHalo7 | 5 |
|  |  | attP40 | - | UAS-LA-Halo2 | 5 |
| Bxb1 | heat shock | attP18 | - | HeatShock-Bxb1 | - |
|  | heat shock | attP40 | - | HeatShock-Bxb1 | - |
|  | heat shock | P element | - | HeatShock-Bxb1 | 4 |
|  | UAS | VK00027 | yes | UAS->FlpSTOP>Bxb1 | - |
|  | UAS | VK00027 | - | UAS->Bxb1 | 4 |
| SNAPf<br>CLIPf<br>Halo | UAS | attP2 | yes | UAS-ChemicalBrainbow | - |
|  |  | VK00005 | yes | UAS-ChemicalBrainbow | - |
|  |  | attP2,<br>VK00005 | yes | UAS-2xChemicalBrainbow | 4 |

**Table S2:** Genotypes of flies used in each figure. Transgenes marked with an * are not required nor have an effect on the experiment.

| Figure | Genotype in figure | Full Genotype |
| --- | --- | --- |
| 1b | Mz19-Gal4 > UAS-CD4::CLIPf, 2nd chr. | ; Mz19-Gal4 / UAS-CD4::CLIPf ; + / Ki |
| 1b | Mz19-Gal4 > UAS-myr::4xCLIPf, attP40 | ; MZ19-Gal4 / UAS-myr::4xCLIPf in attP40 ; |
| 1b | Mz19-Gal4 > UAS-myr::4xCLIPf, VK00005 | ; Mz19-Gal4 / CyO;UAS-myr::4xCLIPf in VK00005 / + |
| 1b | Mz19-Gal4 > UAS-myr::SNAPf, attP40 | ; MZ19-Gal4 / UAS-myr::SNAPf in attP40-5 ; + / Ki |
| 1b | Mz19-Gal4 > UAS-myr::4xSNAPf, attP40 | ; MZ19-Gal4 / UAS-myr::4xSNAPf in attP40 ; + / MKRS |
| 1b | Mz19-Gal4 > UAS-myr::4xSNAPf, VK00005 | ; MZ19-Gal4 / CyO;UAS-myr::4xSNAPf in VK00005 / + |
| 1c | GH146-LexA > LexAop2-myr::4xCLIPf, attP40 | hsFLP* ; GH146-LexA / LexAop2-myr::4xCLIPf in attP40 ; MB247-Gal4*,QUAS-mtdTomato*/MKRS |
| 1c | GH146-LexA > LexAop2-myr::4xSNAPf, attP40 | hsFLP* ; GH146-LexA / LexAop2-myr::4xSNAPf in attP40 ; MB247-Gal4*, QUAS-mtdTomato*/MKRS |
| 1d | Mz19-Gal4 > UAS-myr::4xSNAPf, LexAop2-myr::4xCLIPf | ; MZ19-Gal4 / LexAop2-myr::4xCLIPf in attP40 ; OrcoLexAVP16 / UAS-myr::4xSNAPf in VK00005 |
| 2b | Mz19-Gal4 > UAS-myr::Halo2, attP40 | ; MZ19-Gal4, UAS-myr::Halo2 in attP40/CyO ; TM6B / + |
| 2b | Mz19-Gal4 > UAS-Halo7::CAAX, attP40 | ; MZ19-Gal4 / UAS-Halo7::CAAX in attP40 ; |
| 2b | Mz19-Gal4 > UAS-3xHalo7::CAAX, attP40 | ; MZ19-Gal4 / UAS-3xHalo7::CAAX in attP40 ; |
| 2b | Mz19-Gal4 > UAS-7xHalo7::CAAX, attP40 | ; MZ19-Gal4 / UAS-7xHalo7::CAAX in attP40 ; |
| 3a | IR84a-Gal4 > UAS-mCD8::GFP | ; UAS-mCD8::GFP ; IR84a-Gal4 |
| 3a | IR84a-Gal4 > UAS-myr::4xSNAPf | ; UAS-myr::4xSNAPf in attP40 ; IR84a-Gal4 |
| 3a, b | IR84a-Gal4 > UAS-3xHalo7::CAAX | ; UAS-3xHalo7::CAAX in attP40 ; IR84a-Gal4 |
| 4c.i | MB247-Gal4 | LexAop2-mCD8::GFP*, UAS-mCD8::RFP/+ ; ; MB247-Gal4 / + |
| 4c.ii | MB247-Gal4 ∩ nSyb-LexA | ; LexAop2-myr::>HA-BxbSTOP>4xSNAPf / nSyb-LexAP65 ; UAS>Bxb1 / MB247-Gal4, QUAS-mtdTomato* |
| 4c.iii | Mz19-Gal4 | ; MZ19-Gal4 / UAS-myr::4xCLIPf in attP40 ; |
| 4c.iv | Mz19-Gal4 ∩ nSyb-LexA | ; nSyb-LexAP65<br>LexAop2-myr::>HA-BxbSTOP>4xSNAPf / MZ19-Gal4 ; UAS>Bxb1 / + |
| 4c.v | Mz19-Gal4 ∩ MB247-LexA | ; Mz19-Gal4 /<br>LexAop2-myr::>HA-BxbSTOP>4xSNAPf ;<br>MB247-LexA / UAS-Bxb1 |
| 4d | Fru-Gal4 > 2 x Brainbow cassettes | ; hsBxb1 / CyO ; Fru-Gal4 / UAS-myr::>HA-BxbSTOP>SNAPf>CLIPf>Halo2<br>in attP2, UAS-myr::>HA-BxbSTOP>SNAPf>CLIPf>Halo2 in<br>VK00005 |
| 5a | Mz19-Gal4 > UAS-Syt::Halo7 | Mz19Gal4 / UAS-Syt::Halo7 in attP40 |
| 5a | Mz19-Gal4 > UAS-3xSyt::Halo7 | Mz19Gal4 / UAS-3xSyt::Halo7 in attP40 |
| 5a | Mz19-Gal4 > UAS-7xSyt::Halo7 | Mz19Gal4 / UAS-7xSyt::Halo7 in attP40 |
| 5b | BG57-Gal4 > UAS-LA::Halo2 | UAS-Dicer2 ; UAS-LA::Halo2 in attP40 / + ;<br>BG57Gal4 / + |
| S1a | Mz19-Gal4 > UAS-CD4::CLIPf on 2nd | ; Mz19-Gal4 / UAS-CD4::CLIPf ; |
| S1a | 50A02-Gal4 > UAS-CD4::CLIPf on 2nd | ; UAS-CD4::CLIPf / + ; 50A02-Gal4 / + |
| S1a | 54F05-Gal4 > UAS-CD4::CLIPf on 2nd | ; UAS-CD4::CLIPf / + ; 54F05-Gal4 / + |
| S1a | 59F02-Gal4 > UAS-CD4::CLIPf on 2nd | ; UAS-CD4::CLIPf / + ; 59F02-Gal4 / + |
| S1a | Mz19-Gal4 > UAS-CD4::CLIPf on 3rd | ; Mz19-Gal4 / + ; UAS-CD4::CLIPf / + ; |
| S1a | 50A02-Gal4 > UAS-CD4::CLIPf on 3rd | ; ; UAS-CD4::CLIPf / 50A02-Gal4 |
| S1a | 54F05-Gal4 > UAS-CD4::CLIPf on 3rd | ; ; UAS-CD4::CLIPf / 54F05-Gal4 |
| S1a | 59F02-Gal4 > UAS-CD4::CLIPf on 3rd | ; ; UAS-CD4::CLIPf / 59F02-Gal4 |
| S1a | 50A02-Gal4 > UAS-mCD8::GFP | 50A02-Gal4 / UAS-IVS-mCD8::GFP in attP2 |
| S1a | 54F05-Gal4 > UAS-mCD8::GFP | 54F05-Gal4 / UAS-IVS-mCD8::GFP in attP2 |
| S1a | 59F02-Gal4 > UAS-mCD8::GFP | 59F02-Gal4 / UAS-IVS-mCD8::GFP in attP2 |
| S1b | Mz19-Gal4 > UAS-myr::4xCLIPf in attP40 | ; Mz19-Gal4 / UAS-myr::4xCLIPf in attP40 ; |
| S1b | 50A02-Gal4 > UAS-myr::4xCLIPf in attP40 | ; UAS-myr::4xCLIPf in attP40 / + ;<br>50A02-Gal4 / + |
| S1b | 54F05-Gal4 > UAS-myr::4xCLIPf in attP40 | ; UAS-myr::4xCLIPf in attP40 / + ;<br>54F05-Gal4 / + |
| S1b | 59F02-Gal4 > UAS-myr::4xCLIPf in attP40 | ; UAS-myr::4xCLIPf in attP40 / + ;<br>59F02-Gal4 / + |
| S1b | Mz19-Gal4 > UAS-myr::4xCLIPf in VK00005 | ; Mz19-Gal4 / + ; UAS-myr::4xCLIPf in<br>VK00005 / + |
| S1b | 50A02-Gal4 > UAS-myr::4xCLIPf in<br>VK00005 | ; ; UAS-myr::4xCLIPf in VK00005 /<br>50A02-Gal4 |
| S1b | 54F05-Gal4 > UAS-myr::4xCLIPf in<br>VK00005 | ; ; UAS-myr::4xCLIPf in VK00005 /<br>54F05-Gal4 |
| S1b | 59F02-Gal4 > UAS-myr::4xCLIPf in<br>VK00005 | ; ; UAS-myr::4xCLIPf in VK00005 /<br>59F02-Gal4 |
| S1b | 50A02-Gal4 > UAS-mCD8::GFP | ; ; 50A02-Gal4 / UAS-IVS-mCD8::GFP in<br>attP2 |
| S1b | 54F05-Gal4 > UAS-mCD8::GFP | ; ; 54F05-Gal4 / UAS-IVS-mCD8::GFP in attP2 |
| S1b | 59F02-Gal4 > UAS-mCD8::GFP | ; ; 59F02-Gal4 / UAS-IVS-mCD8::GFP in attP2 |
| S2a | Mz19-Gal4 > UAS-myr::4xSNAPf in attP40 | ; Mz19-Gal4 / UAS-myr::4xSNAPf in attP40 ; |
| S2a | 54F05-Gal4 > UAS-myr::4xSNAPf in attP40 | ; UAS-myr::4xSNAPf in attP40 / + ; 54F05-Gal4 / + |
| S2a | 59F02-Gal4 > UAS-myr::4xSNAPf in attP40 | ; UAS-myr::4xSNAPf in attP40 / + ; 59F02-Gal4 / + |
| S2a | 54F05-Gal4 > UAS-mCD8::GFP | ; ; 54F05-Gal4 / UAS-IVS-mCD8::GFP in attP2 |
| S2a | 59F02-Gal4 > UAS-mCD8::GFP | ; ; 59F02-Gal4 / UAS-IVS-mCD8::GFP in attP2 |
| S2b | Mz19-Gal4 > UAS-myr::4xSNAPf in attP2 | ; Mz19-Gal4 / + ; UAS-myr::4xSNAPf in attP2 / + ; |
| S2b | 54F05-Gal4 > UAS-myr::4xSNAPf in attP2 | ; ; 54F05-Gal4 / UAS-myr::4xSNAPf in attP2 |
| S2b | 59F02-Gal4 > UAS-myr::4xSNAPf in attP2 | ; ; 59F02-Gal4 / UAS-myr::4xSNAPf in attP2 |
| S2b | 54F05-Gal4 > UAS-mCD8::GFP | ; ; 54F05-Gal4 / UAS-IVS-mCD8::GFP in attP2 |
| S2b | 59F02-Gal4 > UAS-mCD8::GFP | ; ; 59F02-Gal4 / UAS-IVS-mCD8::GFP in attP2 |
| S2c | GH146-LexA > LexAop2-myr::4xCLIPf in VK00005 | GH146-LexA / CyO ; LexAop2-myr::4xCLIPf in VK00005 / + |
| S2c | GH146-LexA > LexAop2-myr::4xSNAPf in VK00005 | GH146-LexA / CyO ; LexAop2-myr::4xSNAPf in VK00005 / + |
| S3a | OR22a-Gal4 > UAS-mCD8::GFP | ; OR22a-Gal4 / UAS-mCD8::GFP ; |
| S3a | OR22a-Gal4 > ; OR22a-Gal4 / UAS-mCD8::GFP ; | ; OR22a-Gal4 / ; OR22a-Gal4 / UAS-mCD8::GFP in attP40 ; |

**Figure S1:**
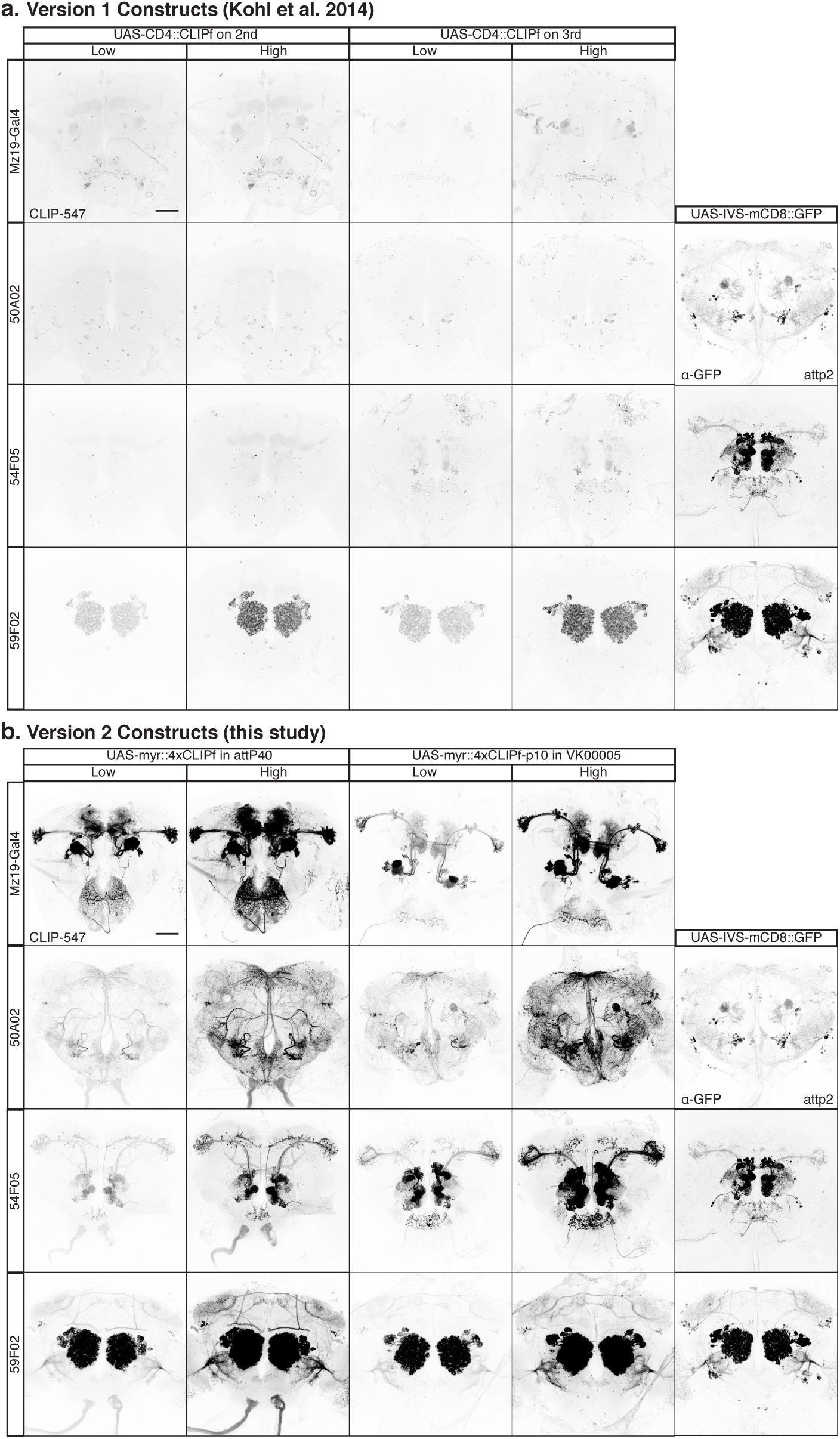
Labeling Gal4 drivers with old and new CLIPf reporters. (**a**) Comparison of UAS-CD4::CLIPf, on 2nd and 3rd chromosomes from Kohl et al. (2014), labeling neurons in the Mz19-Gal4, GMR-50A02-Gal4, GMR-59F02-Gal4 and GMR-54F05-Gal4 expression patterns. The right most panels show the labeling of GMR-50A02-Gal4, GMR-59F02-Gal4 and GMR-54F05-Gal4 neurons using UAS-IVS-mCD8::GFP in attP2. (**b**) Comparison of UAS-myr::4xCLIPf, in attP40 and VK00005, labeling neurons in the Mz19-Gal4, GMR-50A02-Gal4, GMR-59F02-Gal4 and GMR-54F05-Gal4 expression patterns. The right most panels again show the labeling of GMR-50A02-Gal4, GMR-59F02-Gal4 and GMR-54F05-Gal4 neurons using UAS-IVS-mCD8::GFP in attP2. All images of chemical tagging reporters taking using the same confocal settings which achieved non-saturated images with the new transgenes. Right-most panels showing GFP staining are reproduced from http://flweb.janelia.org/cgi-bin/flew.cgi and were published in Jenett et al. (2012). All scale bars are 50 μm.

**Figure S2:**
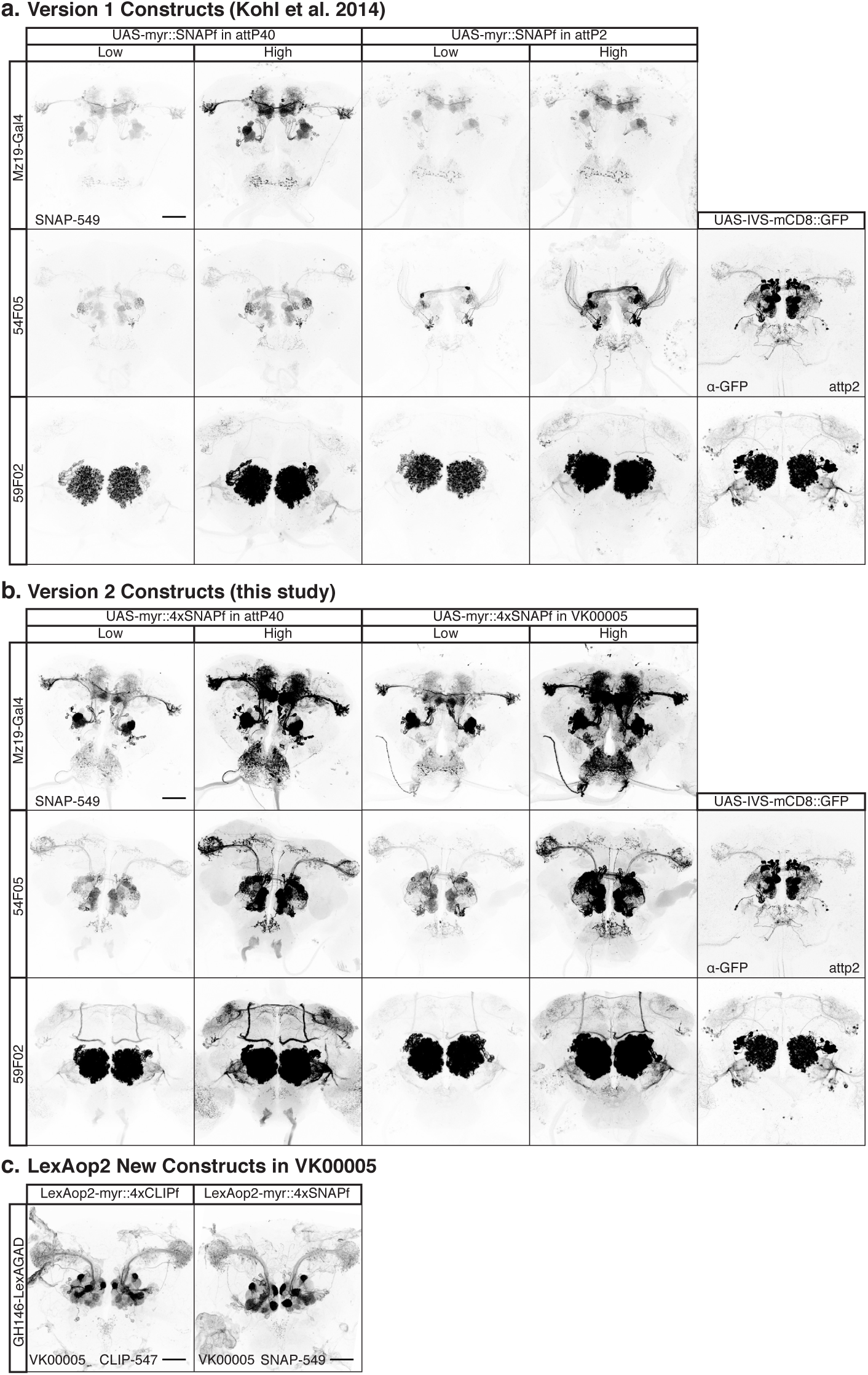
Labeling of Gal4 drivers with old and new SNAPf reporters. (**a**) Comparison of version UAS-myr::SNAPf, in attP40 and attP2 from Kohl et al. (2014), labeling neurons in the Mz19-Gal4, GMR-59F02-Gal4 and GMR-54F05-Gal4 expression patterns. The right most panels show the labeling of GMR-59F02-Gal4 and GMR-54F05-Gal4 neurons using UAS-IVS-mCD8::GFP in attP2. (**b**) Comparison of UAS-myr::4xSNAPf, in attP40 and VK00005, labeling neurons in the Mz19-Gal4, GMR-59F02-Gal4 and GMR-54F05-Gal4 expression patterns. The right most panels again show the labeling of GMR-59F02-Gal4 and GMR-54F05-Gal4 neurons using UAS-IVS-mCD8::GFP in attP2. All images of chemical tagging reporters were aquired using the same confocal settings which achieved non-saturated images with the new transgenes. Right-most panels showing GFP staining are reproduced from http://flweb.janelia.org/cgi-bin/flew.cgi and were published in Jenett et al. (2012). All scale bars are 50 μm.

**Figure S3:**
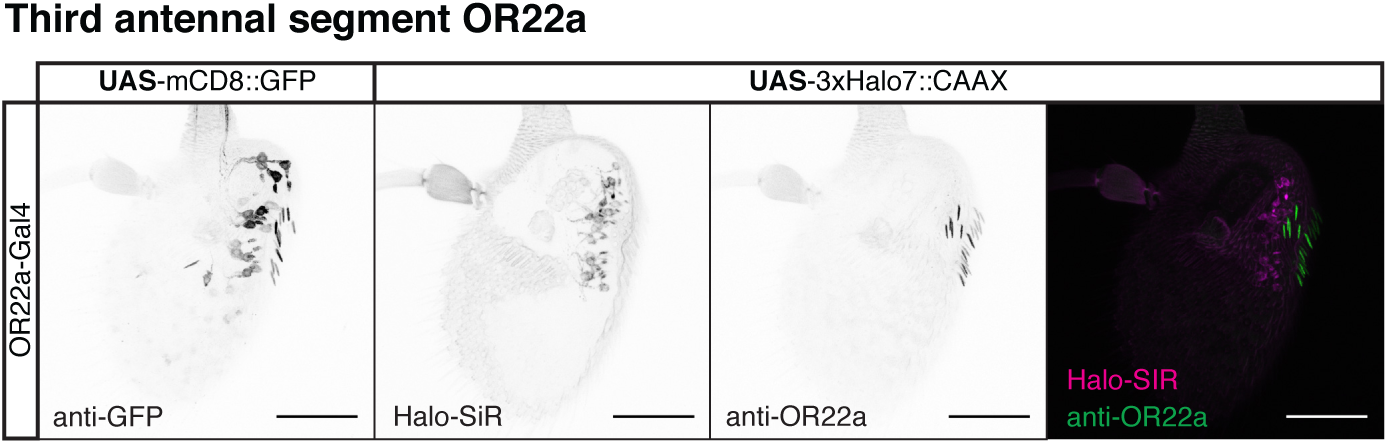
Combining chemical labeling and antibody staining in the antennae. Left panel shows staining of mCD8::GFP in Olfactory Receptor 22a expressing sensory neurons (OR22a). Next three panels show chemical labeling of cell membranes and antibody staining of the OR22a receptor. All panels partial projections of confocal stacks that exclude the cuticle. All inset images are the corresponding confocal full projections. All scale bars are 50 μm.

**Figure S4:**
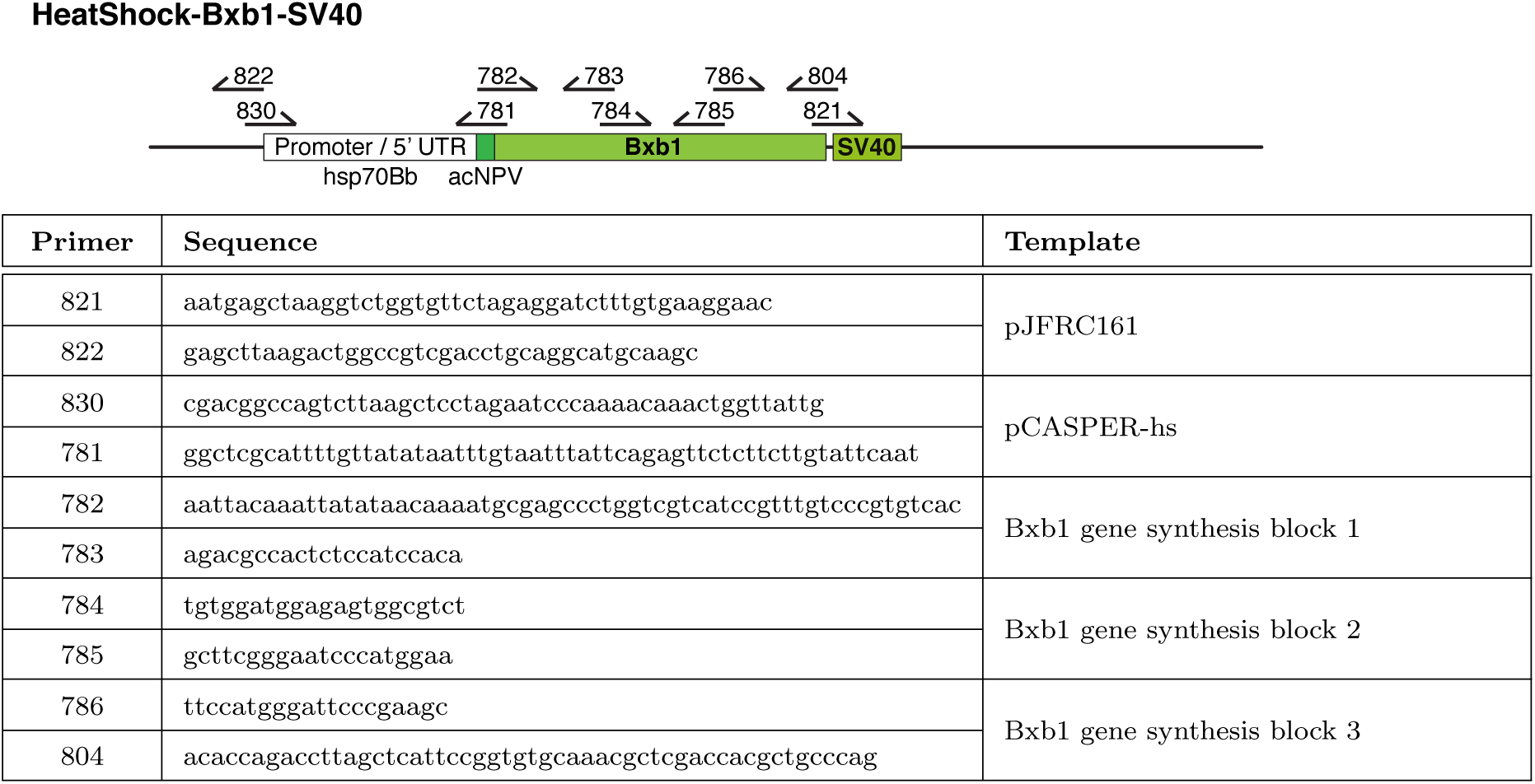
HeatShock-Bxb1-SV40.

**Figure S5:**
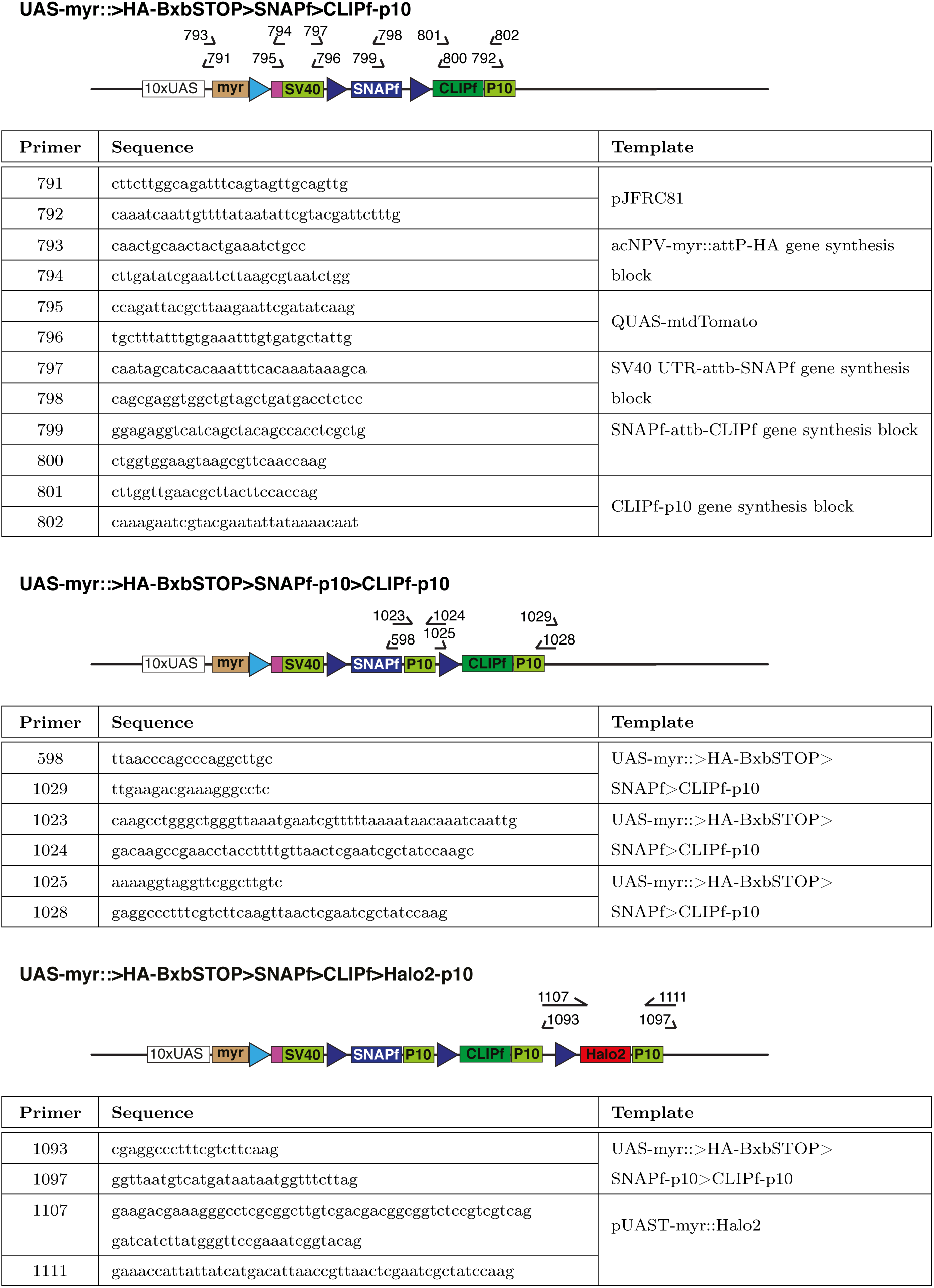
UAS-myr::>HA-BxbSTOP>SNAPf>CLIPf>Halo2.

**Figure S6:**
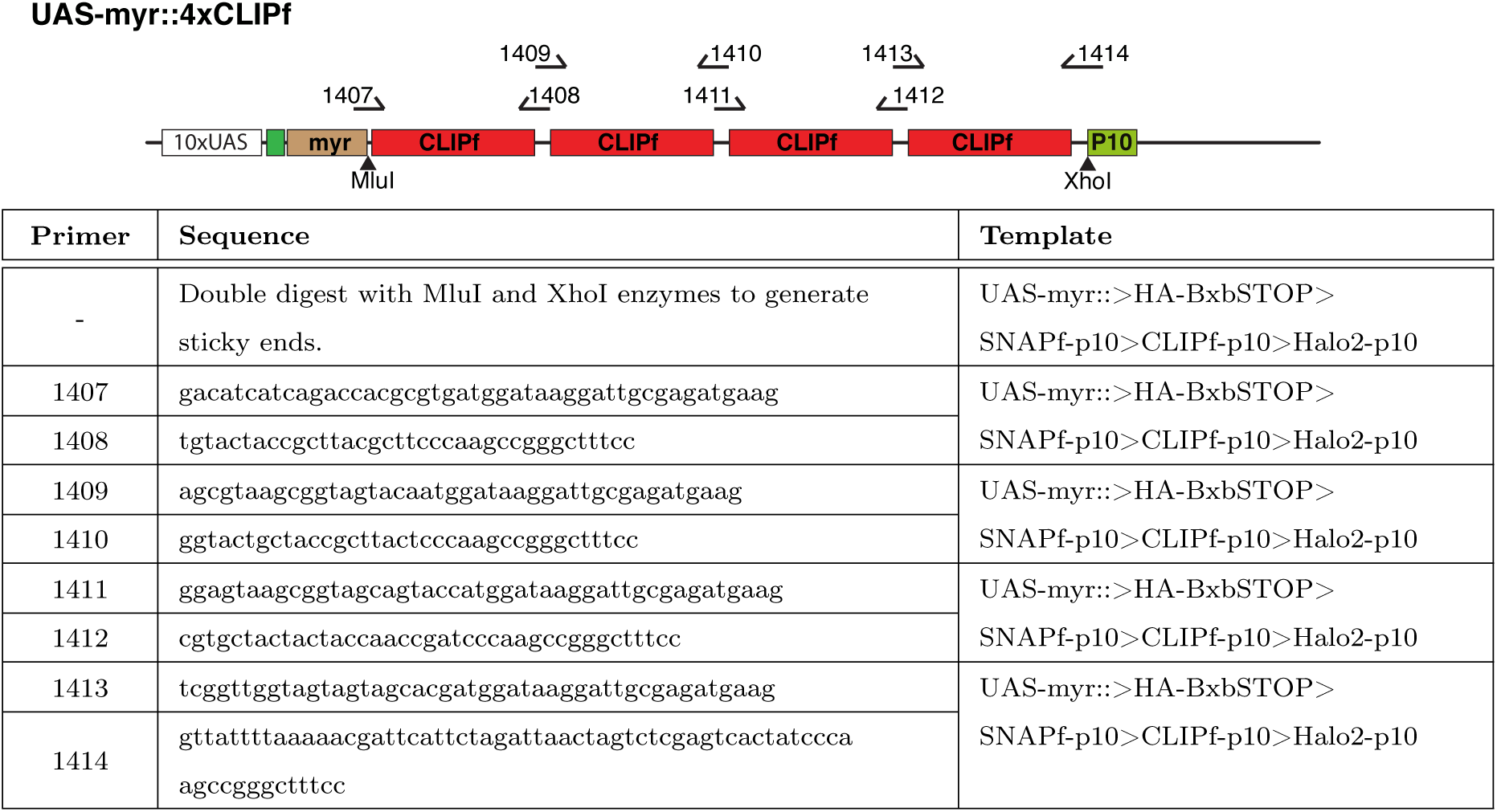
UAS-myr::4xCLIPf.

**Figure S7:**
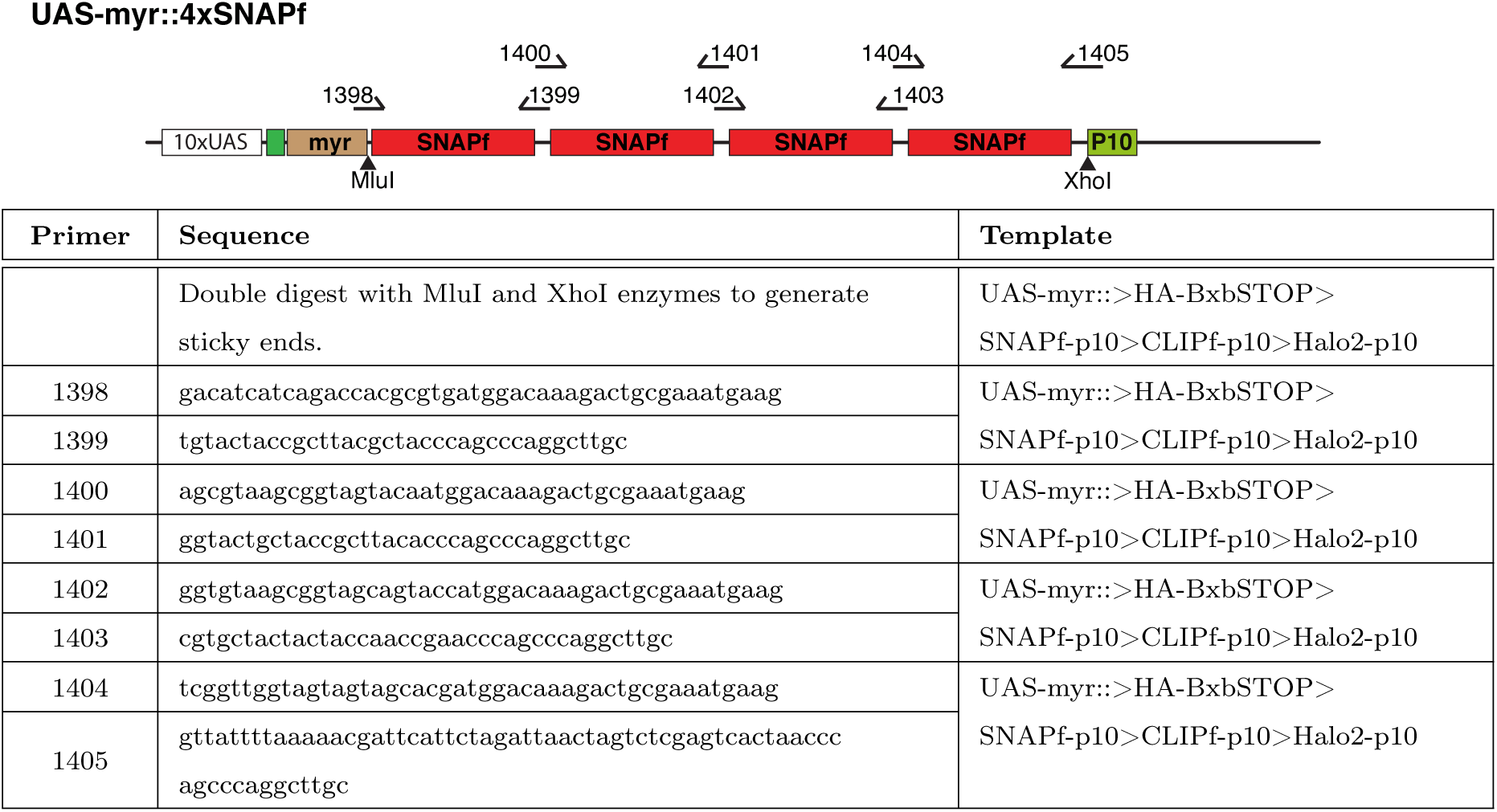
UAS-myr::4xSNAPf.

**Figure S8:**
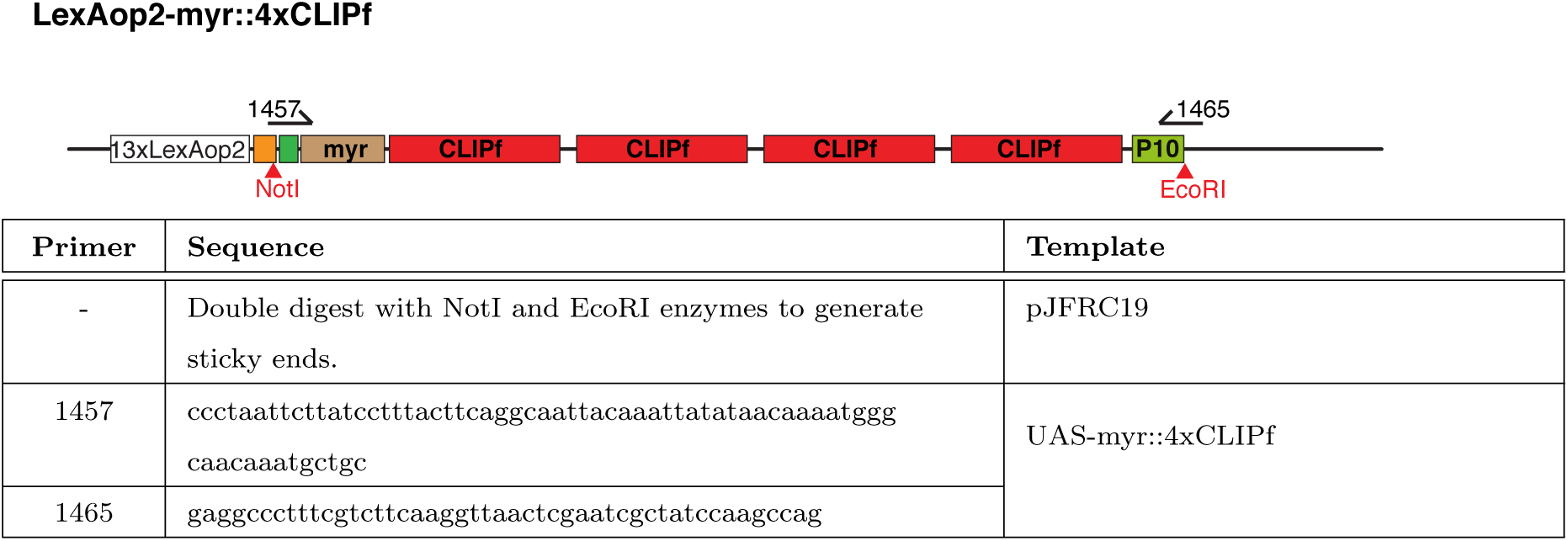
LexAop2-myr::4xCLIPf. Red restriction enzymes indicate that the site is destroyed during the assembly reaction.

**Figure S9:**
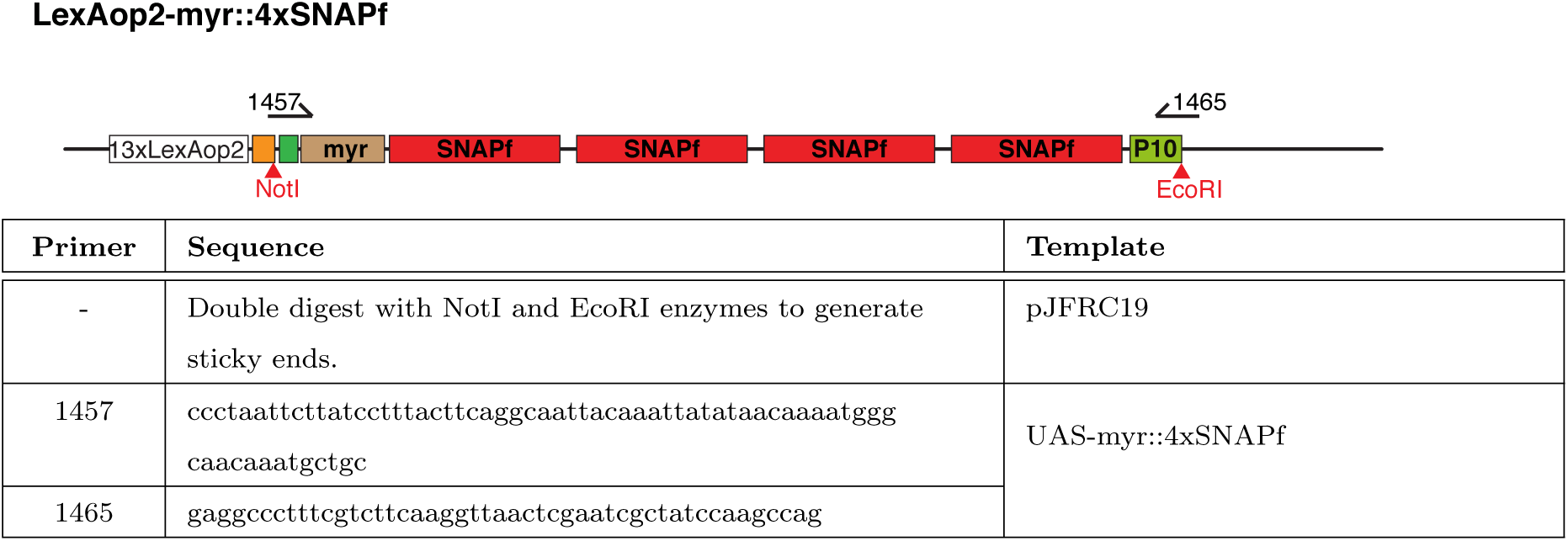
LexAop2-myr::4xSNAPf.

**Figure S10:**
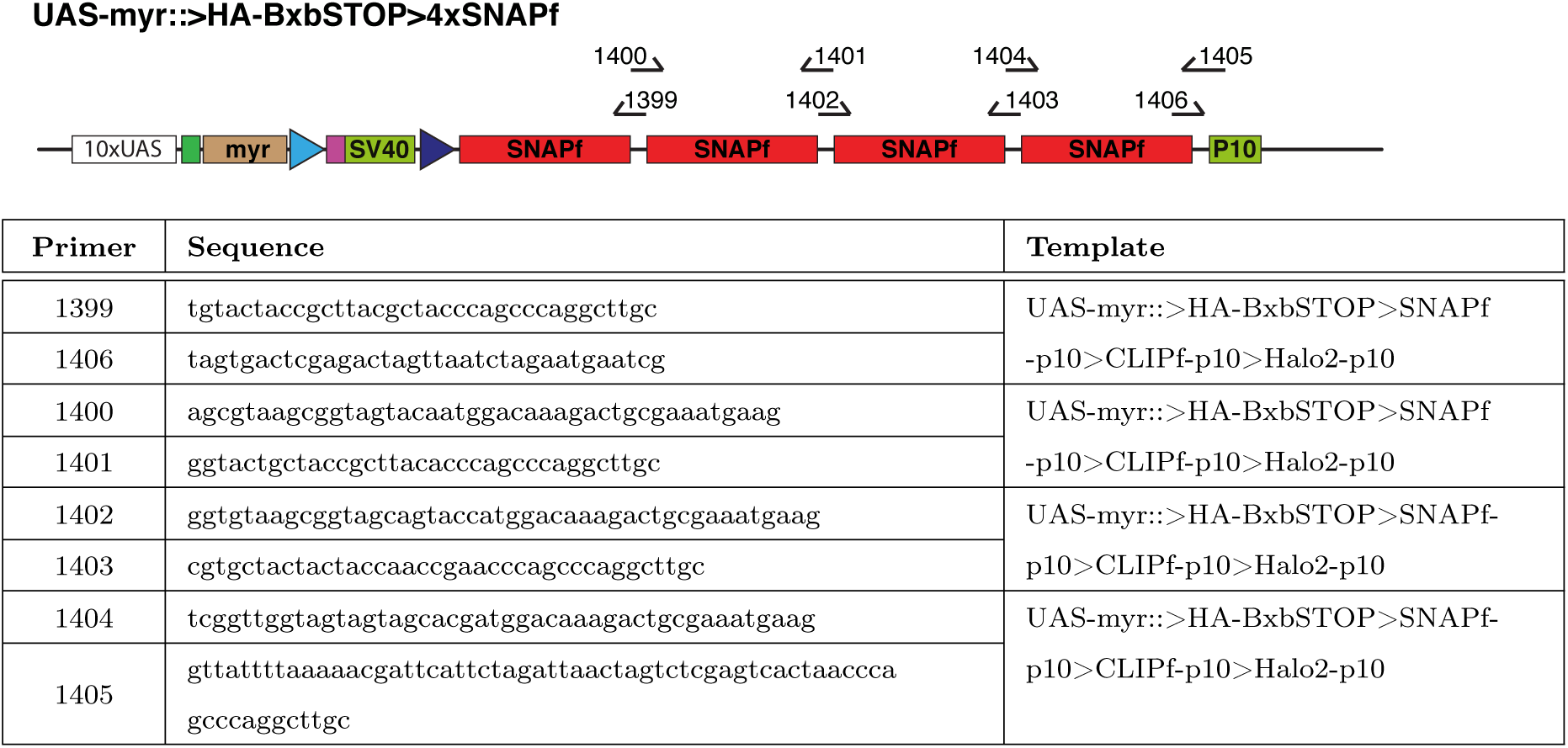
UAS-myr::>HA-BxbSTOP>4xSNAPf.

**Figure S11:**
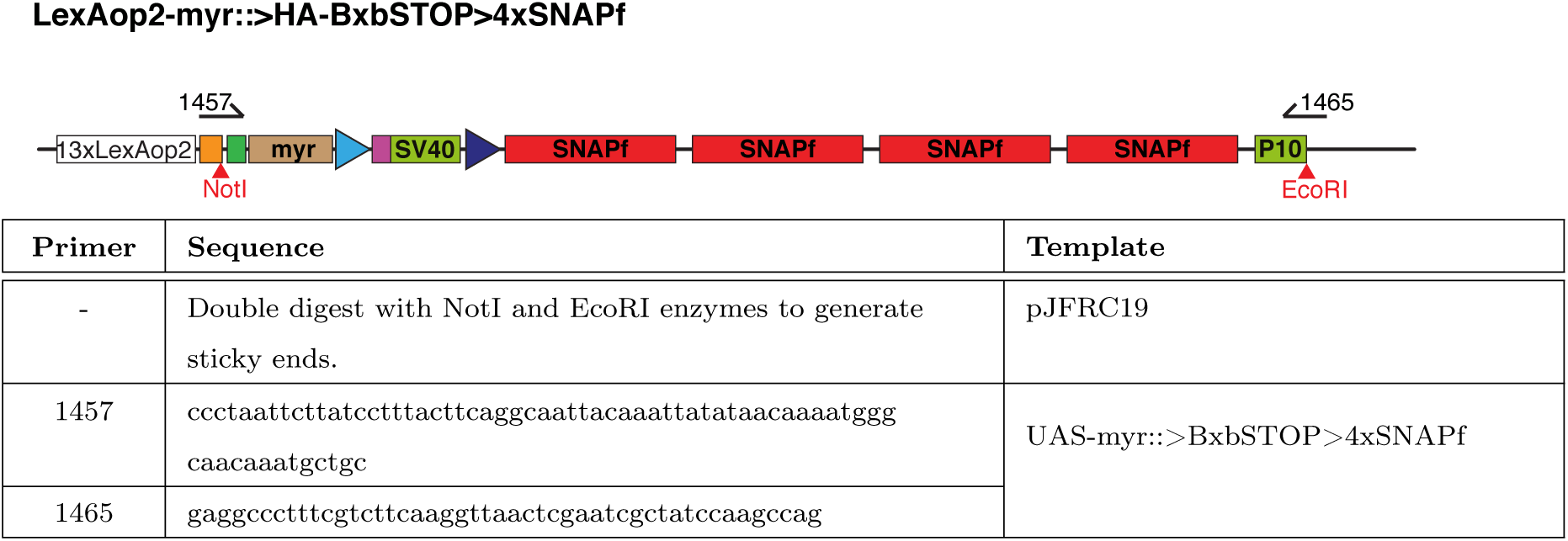
LexAop2-myr::>HA-BxbSTOP>4xSNAPf.

**Figure S12:**
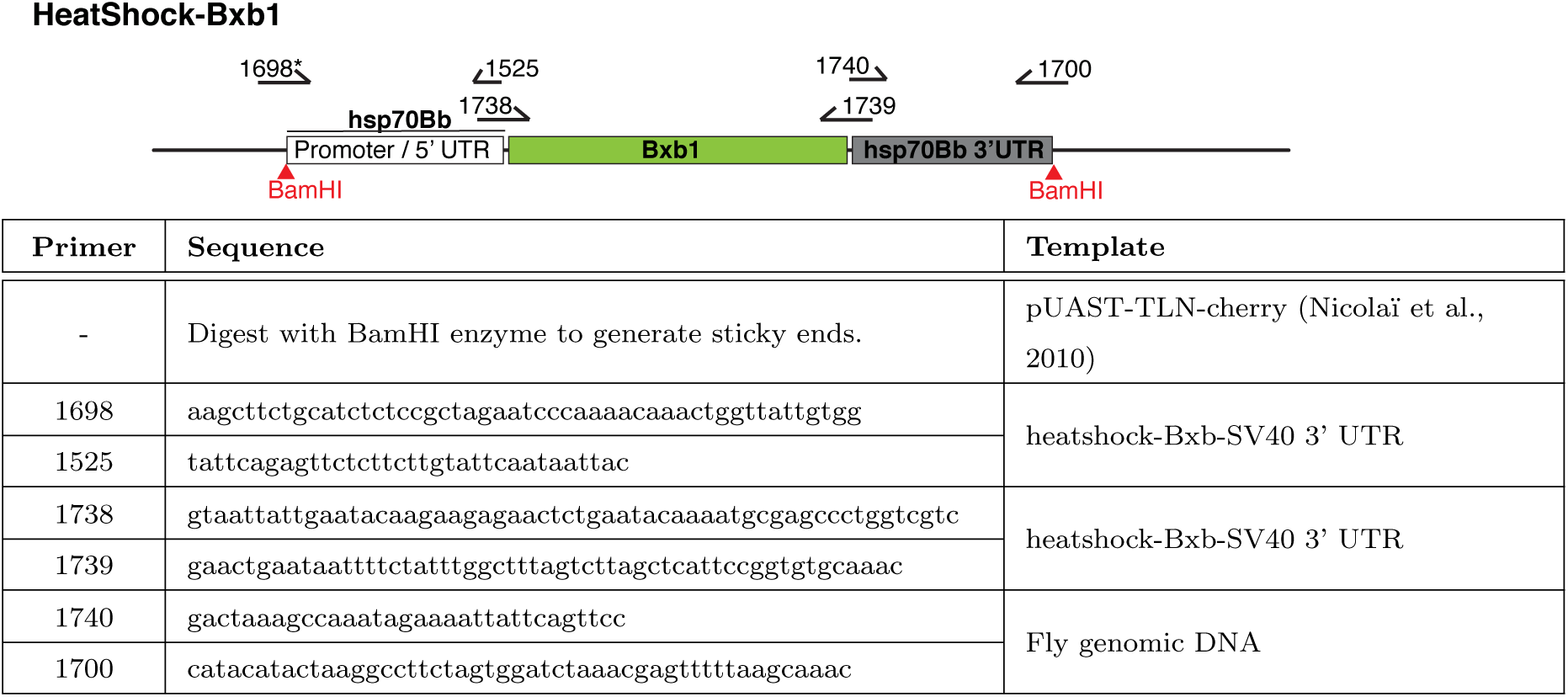
HeatShock-Bxb1. Red restriction enzymes indicate that the site is destroyed during the assembly reaction. Part of the sequence for primer 1698* was not found on the cloned construct; the difference being upstream of the functional sequences does not affect its activity.

**Figure S13:**
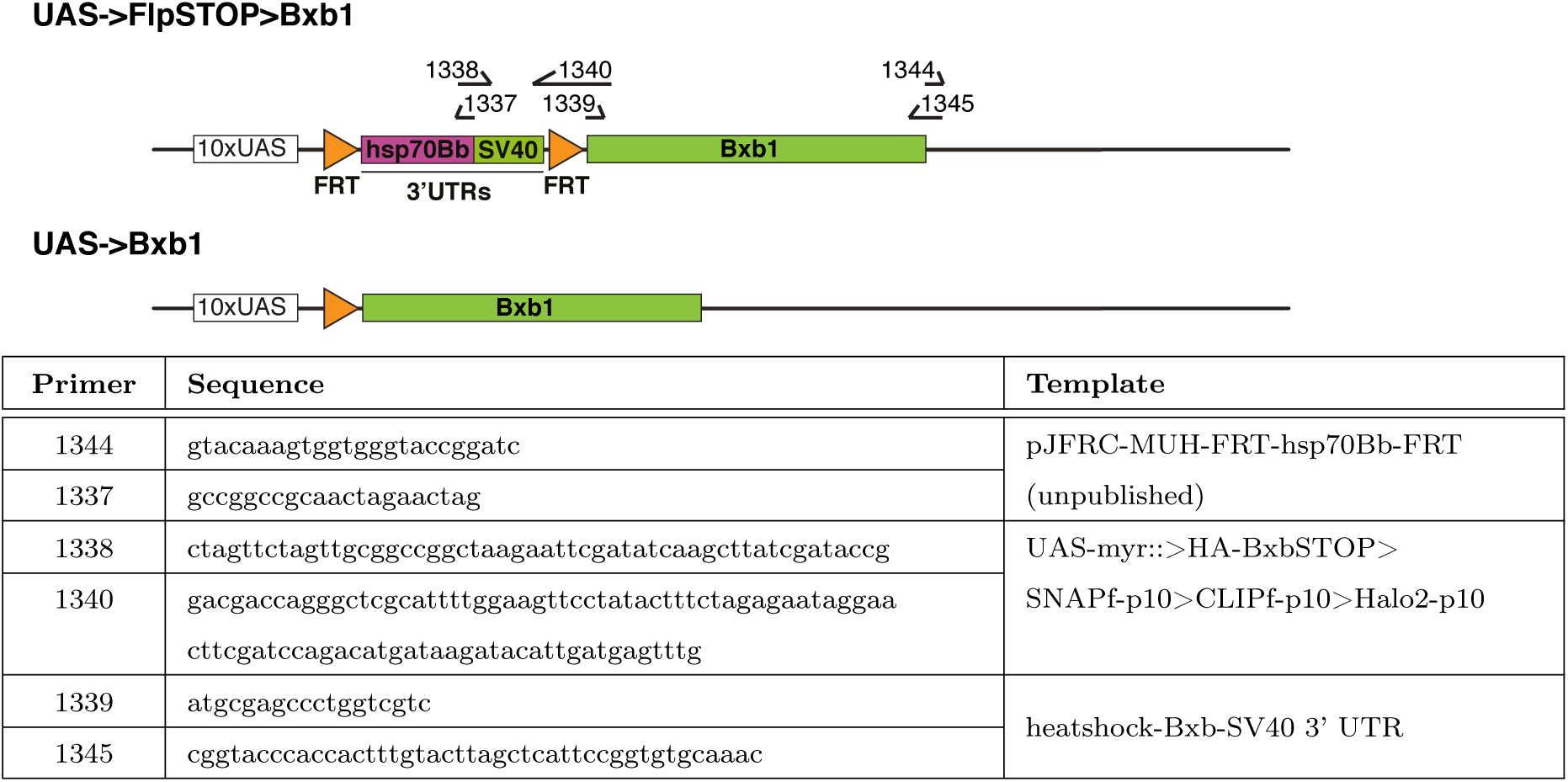
UAS->FlpSTOP>Bxb1. UAS->Bxb1 was obtained by activating UAS-FLP on the germ line of males using nanos-Gal4. Flp recombination induces removal of the stop cassette in the germ line and allowed the establishment of a stock.

**Figure S14:**
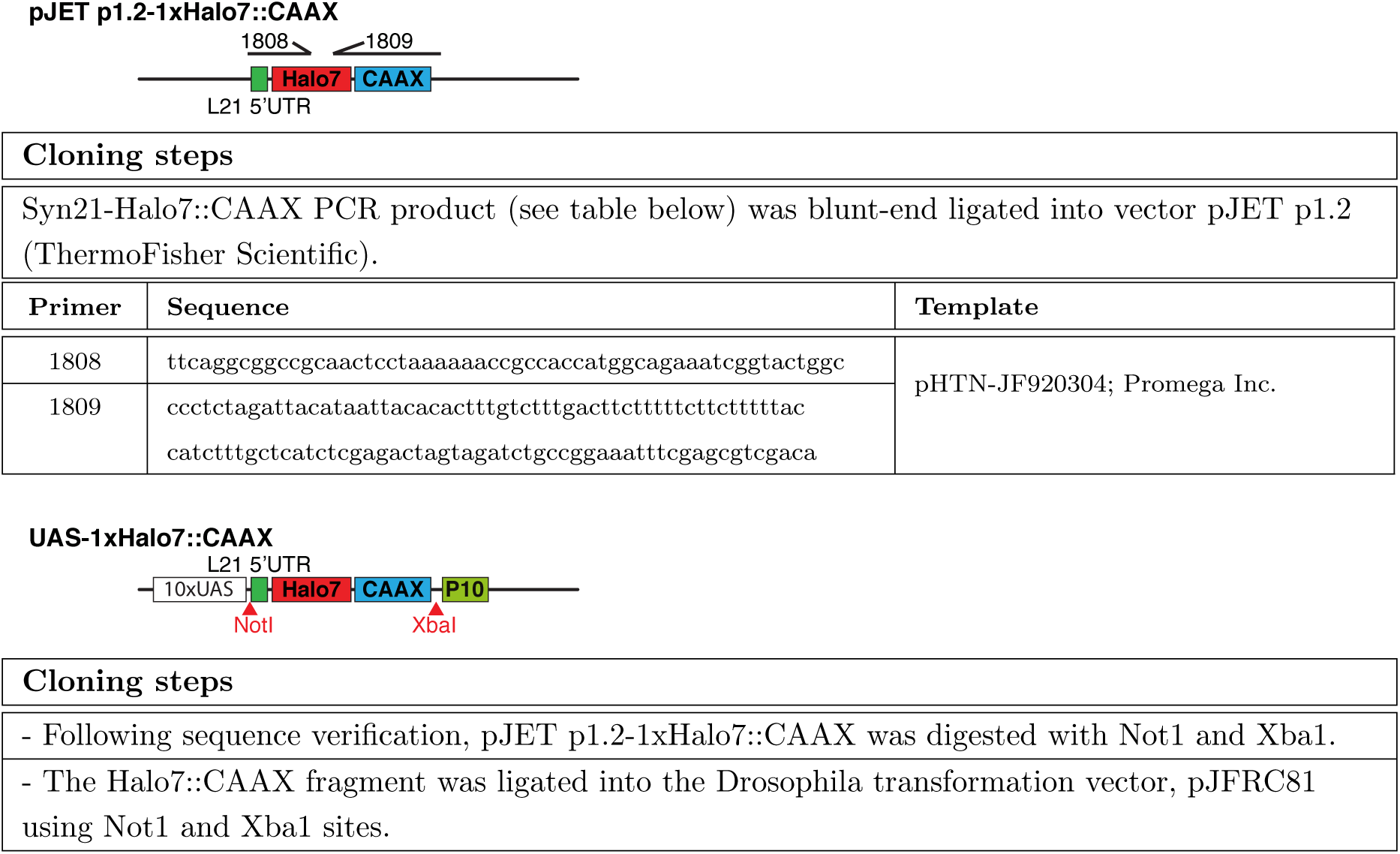
UAS-1xHalo7::CAAX.

**Figure S15:**
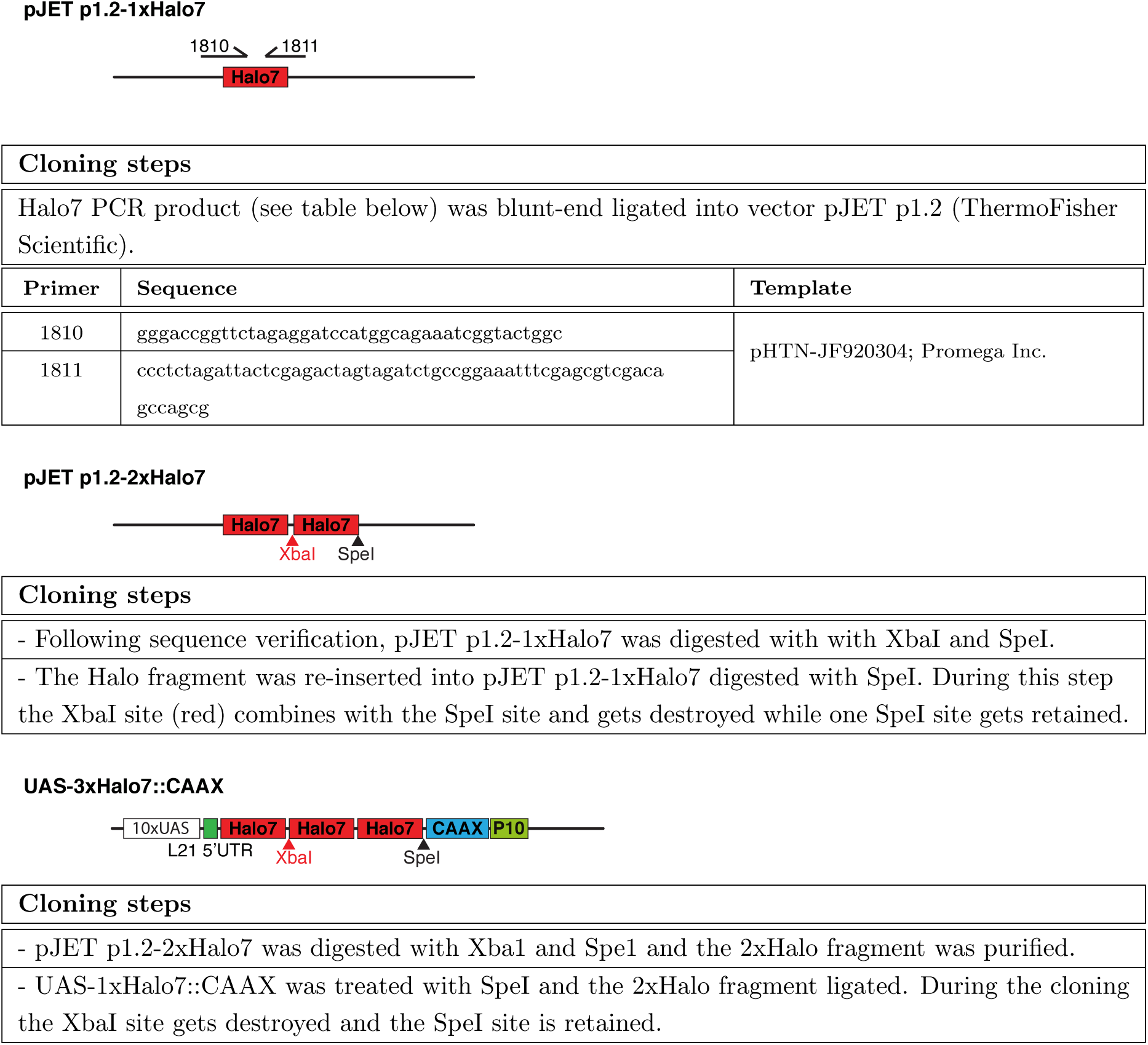
UAS-3xHalo7::CAAX.

**Figure S16:**
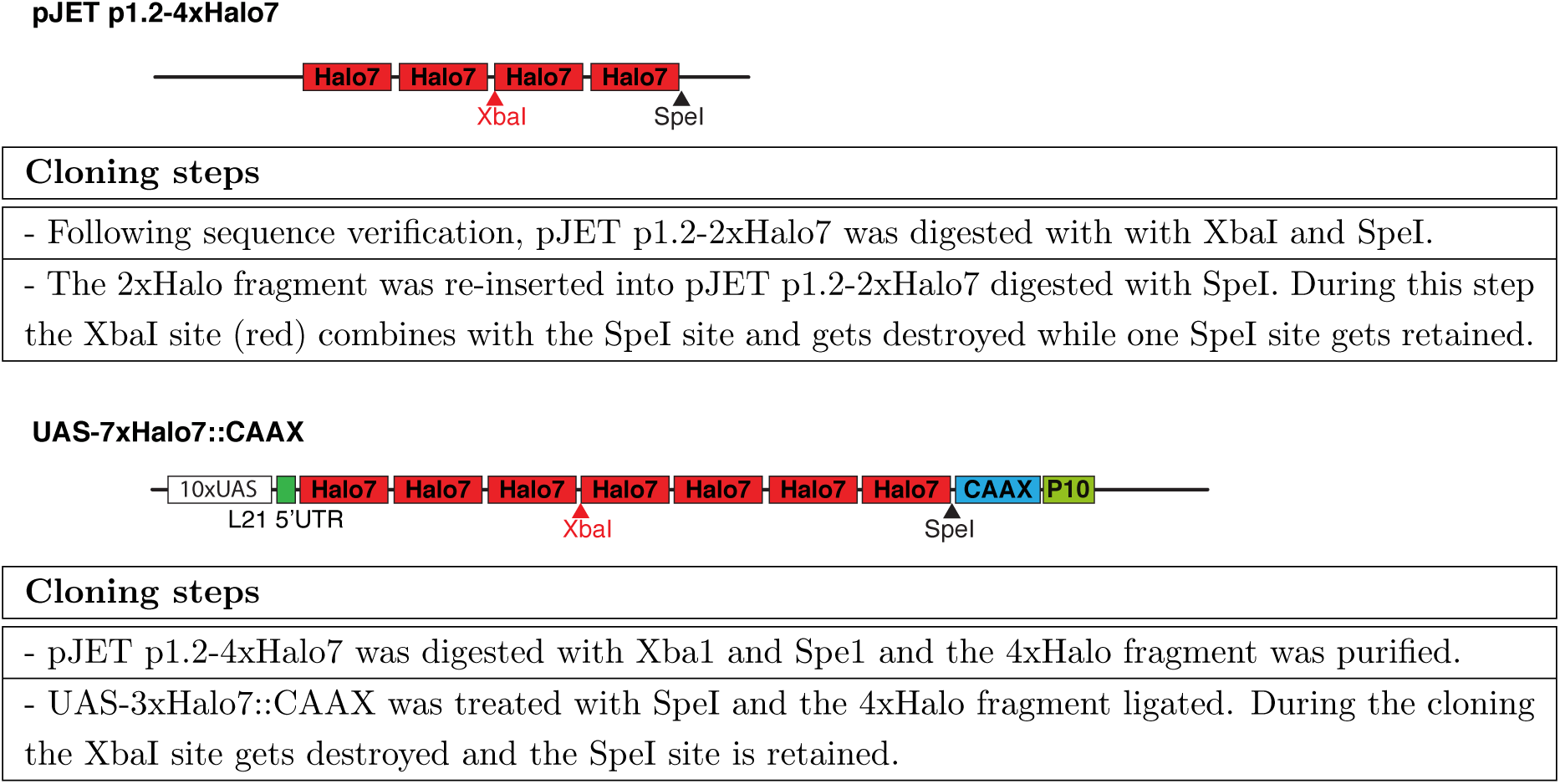
UAS-7xHalo7::CAAX.

**Figure S17:**
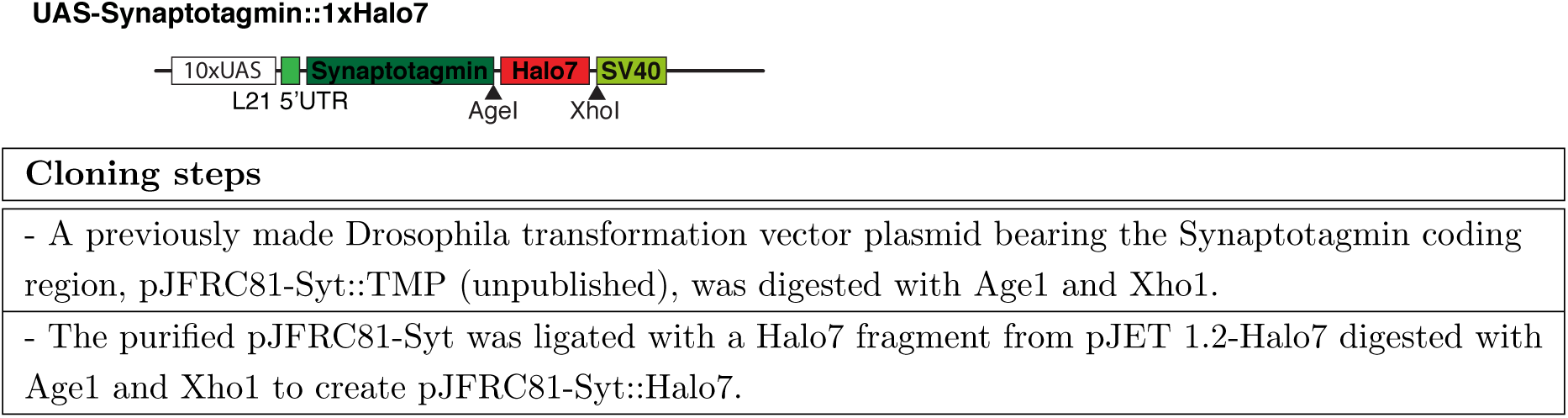
UAS-Synaptotagmin::1xHalo7.

**Figure S18:**
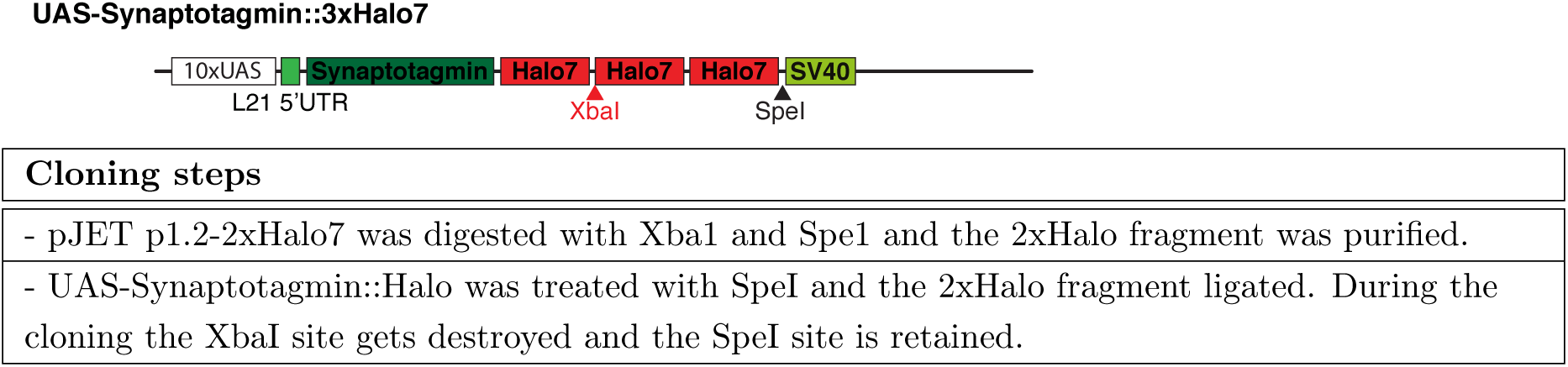
UAS-Synaptotagmin::3xHalo7.

**Figure S19:**
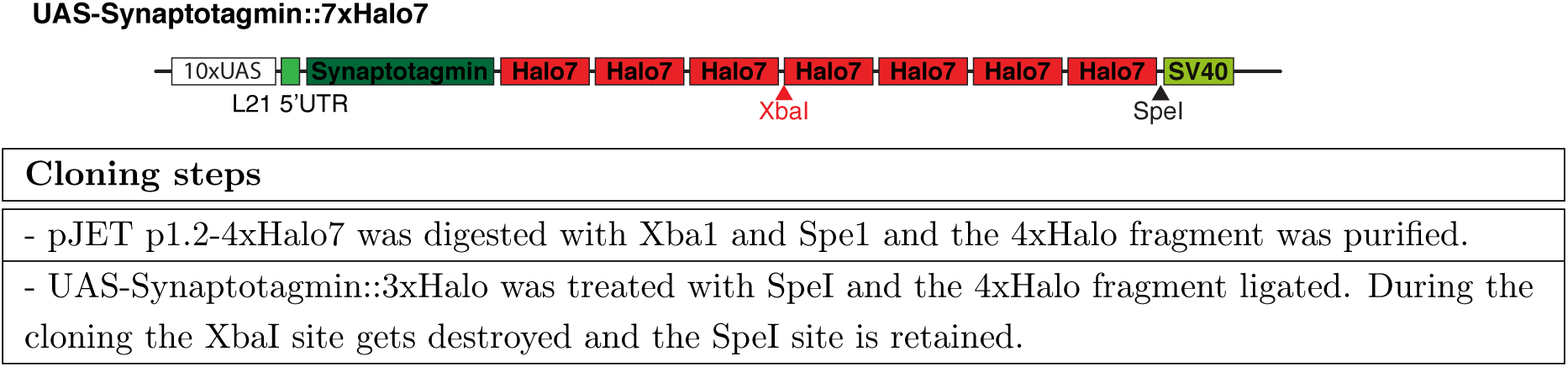
UAS-Synaptotagmin::7xHalo7.

**Figure S20:**
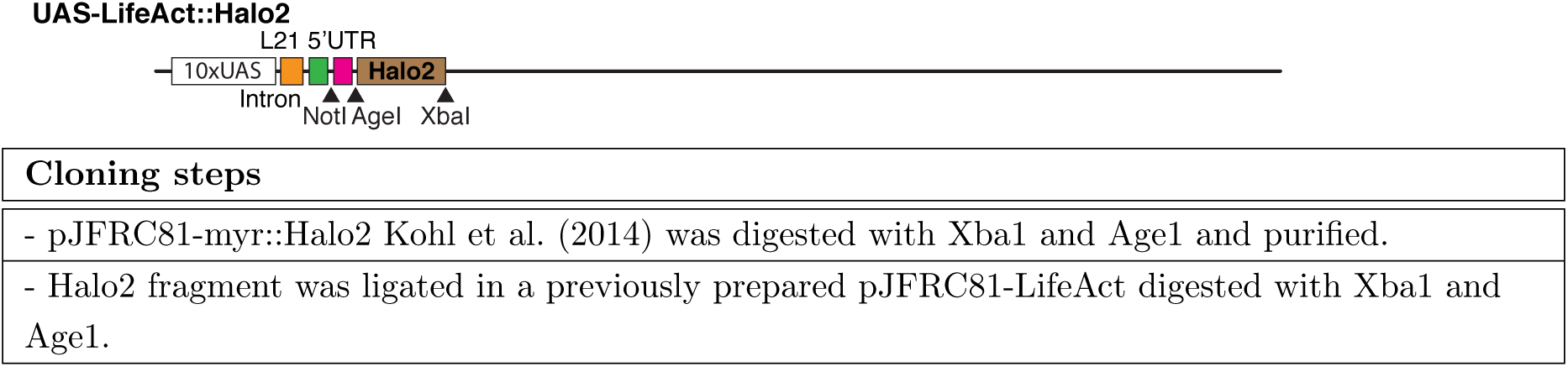
UAS-LA-Halo2.

